# Distinct microbial response to organic matter from nitrogen-starved and virus-infected phytoplankton

**DOI:** 10.1101/2025.08.05.668661

**Authors:** Shira Givati, Eyal Rahav, Osnat Weissberg, Dikla Aharonovich, Noga Barak, Qinglu Zeng, Daniel Sher

## Abstract

When phytoplankton die they release dissolved organic matter (DOM) that feeds co-occurring heterotrophic bacteria. We show that death due to phage infection and nitrogen starvation result in different changes to the macromolecular structure of *Prochlorococcus*, a globally abundant cyanobacterium, and that the resulting DOM elicits different microbial responses. Viral infection led to increased RNA and DNA content in *Prochlorococcus* whereas nitrogen starvation led to a lower protein content. DOM released from phage-infected cells induced high secondary (bacterial) production, while DOM from starved cells increased dark (heterotrophic) primary production in natural microbial communities from the oligotrophic Eastern Mediterranean Sea. Through 16S and 18S amplicon sequencing, metagenomics, and laboratory experiments we identify *Alteromonadaceae* and *Rhodobacteraceae* as heterotrophic taxa responding differently to the two DOM sources. We propose that distinct forms of phytoplankton mortality drive shifts in microbial community metabolism, including differential activity of pathways for heterotrophic carbon fixation, likely through anaplerotic reactions.

## Introduction

Heterotrophic bacteria in the ocean rely on dissolved organic matter (DOM) released by phytoplankton, including cyanobacteria, as a primary resource for growth [1,2]. DOM is released by phytoplankton as they grow (e.g. through exudation or vesicle release, [3]), and as they die, with the specific cause of death potentially affecting the biochemical structure of the phytoplankton and the composition of the released DOM. For example, *Prochlorococcus*, which is the numerically most abundant phytoplankton cell in the oligotrophic oceans [4], contributes ∼8.5% of the net global marine primary productivity estimated at ∼4 Gt C y^-1^ [5]. As *Prochlorococcus* cells die, they release a diverse mixture of organic compounds, thereby altering the complex DOM pool in the water column [6–8]. *Prochlorococcus* die mostly through grazing, phage infection and “other” sources of mortality such as UV damage and, potentially, nutrient starvation, but the relative contribution of each cause of death is currently not well constrained. Estimates range between ∼66-96% for grazing, 4-40% for phage infection and 1-27% for “other” sources of mortality including starvation [9–12]. Given that the pulse of organic carbon (C) released when *Prochlorococcus* cells die could account for more than 75% of the daily production at the surface of the oligotrophic ocean [13], understanding what affects the composition of this photosynthetically fixed C and how it affects the surrounding microbial ecosystem could be of significant biogeochemical and ecological importance.

Different types of mortality, such as starvation and phage infection, can alter the composition of released DOM because each cause of death is associated with distinct changes in cellular physiology, resource allocation and macromolecular structure. *Prochlorococcus* cells experiencing nitrogen starvation in the lab have decreased levels of protein and RNA along with elevated carbon content, potentially from storage carbohydrates or lipids [14,15]. Similar changes are seen also in other phytoplankton [16,17]. In contrast, little is known about how virus or phage infection affects the macromolecular structure of phytoplankton, but since phage infections results in the production of protein- and nucleic-acid rich virions, these pools are expected to increase [18]. Indeed, phage infection has been shown to change host metabolism and biochemical composition (metabolite composition, [19–21]). As the cells die and lyse, the changes in macromolecular structure are expected to result in concomitant changes in the released DOM [22,23].

Once DOM is released from the dying cells, it fuels secondary bacterial production and growth by surrounding heterotrophic bacteria [24]. During exponential growth in the lab, *Prochlorococcus* releases a variety of organic compounds, including highly labile compounds - amino acids and carboxylic acids such as pyruvate, acetate, and formate [6,25–29]. As the cells die and lyse, they release also cell biomass, which is expected to be enriched in larger and more complex macromolecules such as RNA, DNA, large protein complexes, lipids and other storage polymers. *Prochlorococcus* viral lysate was found to contain more labile, bioavailable DOM (e.g., amino acids, protein-like, and low-molecular-weight compounds), promoting greater bacterial growth than *Prochlorococcus* exudate [22]. In addition, each of these different types of macromolecules can fuel the growth of different heterotrophic bacteria [30–33]. Therefore, it is expected that changes in the biochemical structure of the cells due to different forms of mortality could impact the structure and function of surrounding heterotrophic communities. Indeed, previous studies have suggested that phytoplankton cell lysis due to viral infection (including in *Prochlorococcus*) enhances net productivity and organic matter recycling, significantly more than in ecosystems with low viral infection [22,34–37]. In the ocean, this process is known as the “viral shunt”, which plays a crucial role in oceanic biogeochemical cycles by influencing the microbial loop and facilitating the recycling of organic carbon within the marine environment.

Here, we directly compare how nitrogen starvation and phage infection affect cell quotas of protein, RNA and DNA in *Prochlorococcus* (Figure 1A). We chose to measure these major macromolecular pools as they are the main constituents of cell biomass, determining the elemental ratio of the cells [40,41]. These macromolecules are also emerging as important variables in phytoplankton models from the cellular to ocean scales [15,42]. We then amended surface seawater from the oligotrophic Eastern Mediterranean Sea (EMS) with DOM derived either from nitrogen-starved *Prochlorococcus* (hereafter referred to as treatment nSM) or viral lysate (treatment vSM) (Figure 1A). As expected, we observed increased secondary production due to vSM, but surprisingly we also observed increased dark carbon fixation (DCF) in response to nSM. To investigate the microbial taxa driving this response (i.e., fix CO_2_ in response to nSM), we employed 16S and 18S amplicon sequencing, metagenomic analyses and targeted experiments using specific bacterial isolates. Ultimately, by dissecting the differential release of organic compounds under starvation and phage infection, we aim to gain a deeper understanding of *Prochlorococcus*’s role in marine carbon cycling and its complex interactions within ambient marine microbial ecosystem.

**Figure 1.**
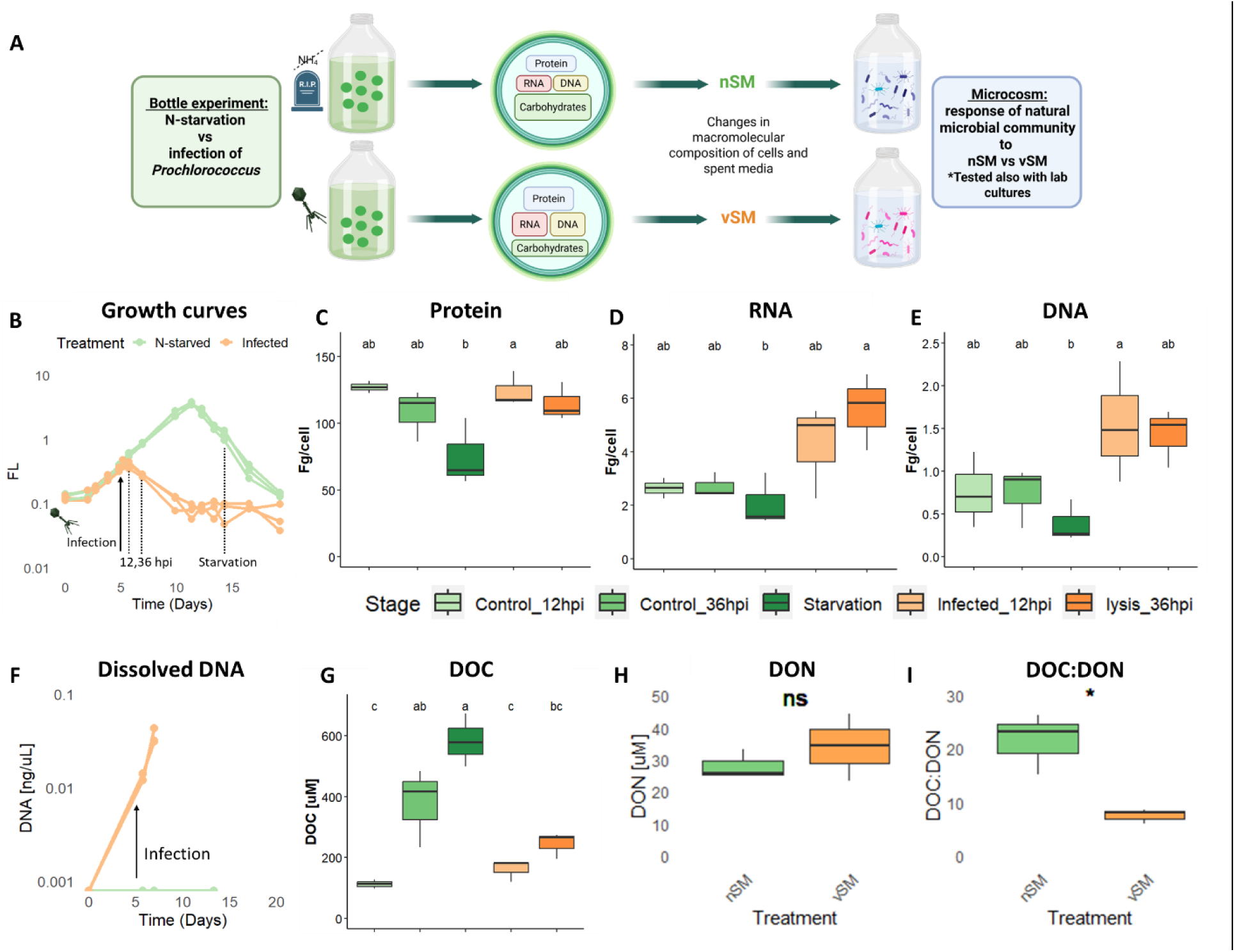
Effect of nitrogen starvation and phage infection on the macromolecular composition of *Prochlorococcus* MED4. (A) Illustration of the experimental set-up. (B) Growth curves of N-starved (green) and infected (orange) *Prochlorococcus* – fluorescence, a proxy for cells numbers. (C) Protein quotas in *Prochlorococcus* cells. (D) RNA quotas. (E) DNA quotas. (F) Dissolved DNA nSM (green) and vSM (orange). (G) Dissolved organic carbon (DOC) concentrations. (H) Dissolved organic nitrogen (DON) in nSM and vSM. (I) DOC:DON ratio in nSM and vSM. Box-Whisker plots show the interquartile range (25th–75th percentile) of the data set. The horizontal line within the box represents the median value (N=3). The different letters above the box plots (C, D, E, G) indicate statistically significant differences among the treatments (one-way ANOVA and post-hoc Tukey test, *p* < 0.05). Asterisks above the box plots (H, I) indicate statistically significant differences between treatments based on t-tests (*p* < 0.05). Abbreviations: hpi = hours post infection; nSM-spent media following nitrogen starvation; vSM - viral spent media.

## Results and discussion

### Phage infection and nitrogen starvation differently affect Prochlorococcus macromolecular composition

We first compared how phage infection and nitrogen starvation influence the macromolecular composition of *Prochlorococcus* MED4 (Figure 1). Cultures were infected by myovirus P-HM2 during early to mid-logarithmic growth, with samples collected after 12 hours post infection (hpi, after one round of infection and lysis [19]), and after 36 hpi when most of the culture has been lysed (likely after several rounds of infections) (Figure 1B). The uninfected cultures were sampled at the same time points, as well as during mid-decline, following the onset of nitrogen starvation (day 14, Figure S1C [43]). Protein quotas per cell, and to lesser extent those of RNA and DNA, decreased in the N-starved *Prochlorococcus* (Figure 1C-E), consistent with prior observations [14,15]. In contrast, RNA and DNA quotas increased for the infected *Prochlorococcus,* and protein quotas remained unchanged (Figure 1C-E).

The increase in intracellular DNA is consistent with synthesis of phage genomic DNA, since P-HM2 has a double-stranded DNA genome [44], but the increase in cellular RNA is more perplexing. In most cells, ribosomes contain ∼80-90% of the total RNA [45], and if this is true in this case it would suggest that the number of ribosomes remains similar or even increases, despite phage infection shutting down major aspects of cell physiology [46]. A previous study of P-HM2 infection in *Prochlorococcus* MED4 showed a decline in host transcripts related to translation and ribosomal proteins, with phage-derived mRNA levels exceed those of the host genes [47]. Some phages encode ribosomal proteins such as bS21 and bL12, which could potentially help stabilize the host ribosomes and/or increase their production [48,49], but no such proteins have been described in P-HM2. It is possible that P-HM2 encodes other proteins or RNAs that stabilize the ribosome. Alternatively, it is possible that most of the measured RNA is in fact phage RNA, and in this case it is possible that ribosomes are in fact depleted, as observed in SAR11, where a substantial fraction (7–15%) of infected cells were found to be ‘ribosome-deprived zombie cells’ [50]. Even if this is true, phage P-HM2 is still able to tightly control the metabolism of its hosts, including potentially protein translation rates, thus avoiding mispackaging of new virions with host rather than viral DNA [47,51].

The DOM released following cell lysis due to phage infection (“viral spent media”, vSM) and nitrogen starvation (“nitrogen-starved spent media”, nSM) also differed. Dissolved organic carbon (DOC) was approximately 2 to 3 times higher in the spent media from nitrogen-starved *Prochlorococcus* (nSM) compared to that from infected *Prochlorococcus* (vSM) (500–650 µmol C L^-1^ versus 200–250 µmol C L^-1^, respectively, Figure 1G). In contrast, dissolved organic nitrogen (DON) concentrations were similar, leading to significantly (*P*-value<0.05, t-test) higher C:N ratio (DOC:DON) in the nSM (Figure 1H, I). This is consistent with previous studies suggesting high DOC release from nitrogen-starved *Prochlorococcus*, potentially due to “overflow metabolism” as the cells continue to fix carbon without being able to form new biomass, thus releasing carbon-rich molecules [7,8,26,52,53]. In contrast, dissolved DNA was measurable only in the vSM (Figure 1F), likely representing released phage particles. Indeed, the vSM contained infective phages (Figure S2).

### DOM from N-starved and infected Prochlorococcus has distinct effects on bacterial activity and dark carbon fixation in natural microbial communities

We next added nSM and vSM to natural microbial communities from the Eastern Mediterranean Sea (EMS) in two separate experiments, one during autumn, when the water column was highly stratified and oligotrophic (Figure S3A-C), and another during spring, when the water column was mixed and nutrients concentrations were much higher (Figure S3D-F). The amendments contained equal amounts of DOC (25 μM, a concentration used in previous studies [30,54]), as well as NH_4_^+^ and PO_4_^2-^ to make sure the microbial communities were not nutrient-limited and could thus respond to the potentially different compositions of organic carbon sources.

During both autumn and spring, the addition of vSM led to a significant (*P*-value<0.05, ANOVA) increase in total cell counts compared to the control where only inorganic nutrients were added, whereas the addition of nSM did not result in a significant change (Figure 2A, Figure S4A). Further examination of the flow cytometry scattergrams indicated that the differences between the nSM and vSM in both experiments were primarily due to a higher increase in the number of low nucleic acid (LNA) cells in response to vSM (*P*-value<0.01, ANOVA, Figure S4C). LNA cells, characterized by reduced fluorescence intensity, small cell size, and compact genomes, are highly abundant in oligotrophic oceans, yet their precise identity remains uncertain [55]. In this study, the LNA population exhibited a larger increase in the intensity of Sybr-green staining in the vSM (*P*-value<0.001, ANOVA, Figure S4F, G). Assuming that the intensity of Sybr-green staining is related to the nucleic acid content of the cells, this might suggest that LNA cells increased both in abundance and activity [56], primarily in response to vSM. Bacterial production (BP), measured as the incorporation of radioactive leucine, also increased significantly during spring in response to the vSM than the nSM (*P*-value<0.001, ANOVA, Figure S5D, BP measurements during autumn were unreliable). Taken together, these results, including increased total cell counts, higher abundance and activity of LNA cells, and elevated bacterial production, suggest that viral lysate elicits faster growth and more heterotrophic activity compared to nSM. This is consistent with experiments performed in the South China Sea with the lysates of P-HM2 and two other phages compared with spent media from exponentially-growing *Prochlorococcus* [22].

**Figure 2.**
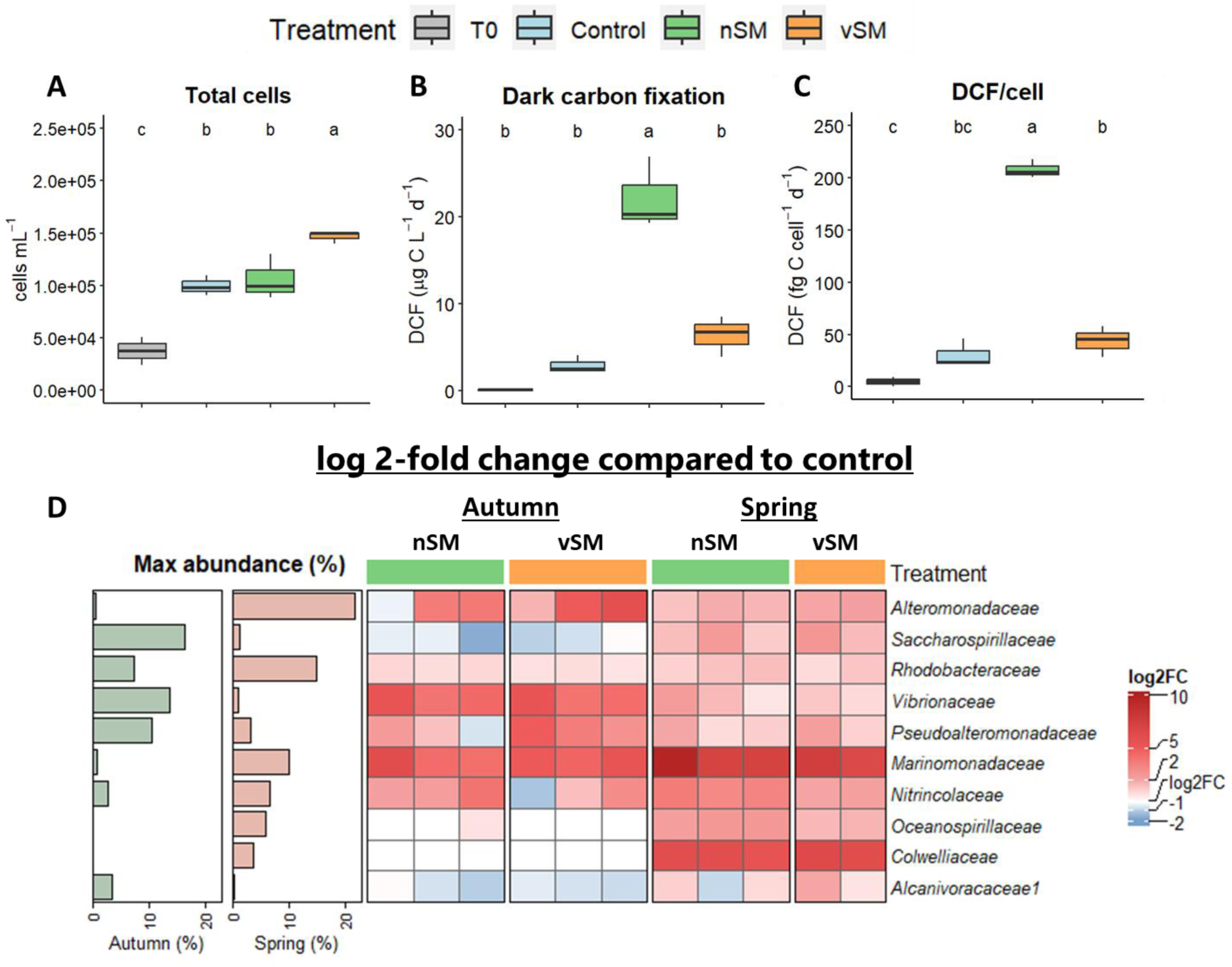
Spent media from N-starved and infected *Prochlorococcus* differently affect natural microbial community activity and structure. (A) Total cells count. (B) Dark inorganic carbon fixation (DCF). (C) Per-cell DCF. Box-Whisker plots show the interquartile range (25th–75th percentile) of the data set. The horizontal line within the box represents the median value (N=3). The different letters above the box plot indicate statistically significant differences among the treatments (one-way ANOVA and post-hoc Tukey test, *p* < 0.05). (D) Heatmap showing the log₂ fold change in relative abundance of the top bacterial families enriched in the nSM and vSM compared to the control, based on 16S rRNA sequencing. Each column is an experimental replicate, and each row is a family for which at least one replicate showed a relative abundance above 1% and a fold change greater than 2 compared to the nutrient-only control. The bar-plots on the left of the heatmap show maximum relative abundance in nSM or vSM in the autumn (green) and spring (pink). For the full bar-plot of the community structure and more discussion please see Figure S6 and Supplementary text 2. Abbreviation: DCF – dark carbon fixation; nSM-spent media following nitrogen starvation; vSM - viral spent media.

In addition to BP, we also measured dark carbon fixation (DCF), the incorporation of CO_2_ into biomass without light energy. DCF can be performed by chemoautotrophs, which use inorganic electron donors such as CO_2_ or H_2_, or by heterotrophic bacteria which use organic matter as a source of energy (see details below). Heterotrophic CO_2_ fixation is increasingly recognized as a significant and previously overlooked contributor to CO_2_ assimilation in marine environments, occurring not only in the deep, dark ocean but also in shallower waters [33,57–62]. Intriguingly, we observed a significant increase in DCF in the nSM but not vSM during both autumn (*P*-value<0.001, ANOVA, Figure 2B, C) and spring (*P*-value<0.01, ANOVA, Figure S5B, C) experiments, reaching in our experimental incubations levels comparable to photosynthesis in the oligotrophic Mediterranean and Red Seas (see Supplementary text 1).

### 16S amplicon sequencing reveals no specific clades enriched in the nSM that could be associated with DCF

We expected the differences in the functional response to nSM and vSM to be mirrored by differences in the community structure, as assessed by amplicon sequencing of the 16S rRNA (which is affected by both relative organism abundance and activity, i.e., number of ribosomes). Specifically, we expected to identify organisms (represented by Amplicon Sequence Variants, ASVs) which exhibit a higher increase in relative abundance in response to nSM than to vSM compared to the no-DOM control. Such ASVs would be good candidate organisms responsible for the increased DCF. However, while the nSM and vSM differed from the nutrient-only controls in both autumn and spring experiments (Figure S6), no ASVs were specifically enriched in the nSM (Figure 2D). Several bacterial families increased in relative abundance in both treatments compared to the controls, including (during autumn) *Vibrionaceae*, *Pseudoalteromonadaceae* and *Rhodobacteraceae* (Figure 2D, Figure S7). During spring, *Alteromonadaceae* rather than *Pseudoalteromonadaceae* increased significantly in both treatments (Figure 2D). These results are in agreement with previous experiments where spent media from *Prochlorococcus* was added to natural communities [22,54,63].

During the Autumn experiment we sequenced also 16S rDNA amplicons (Figure S8), which reflects relative abundance and not activity. *Pseudoalteromonadaceae* and *Vibrionaceae* increased significantly in relative abundance in both rDNA and rRNA amplicons, suggesting rapid growth in response to nSM and vSM. In contrast, the relative abundance of *Rhodobacteraceae* increased only in the rRNA and not the rDNA, suggesting increased activity without necessarily a concomitant increase in cell numbers. The specific *Vibrionaceae* and *Rhodobacteraceae* ASVs responding during autumn and spring were not identical, suggesting temporal differences in the fine-scale community structure or function between the two seasons (Figure S6). Analysis of 18S rRNA sequencing from the autumn experiment revealed no significant changes in eukaryotic community structure (Figure S9).

### Identifying clades with the capacity for DCF using metagenomics

Since the 16S and 18S amplicon sequencing was not able to highlight microbial clades responsible for the increased DCF, we next searched for relevant pathways (and, subsequently, individual genes) in metagenomes from the nSM, vSM and control incubations. Broadly speaking, *autotrophic* CO_2_ fixation is mediated by pathways such as the reverse tricarboxylic acid (rTCA) cycle or the Wood-Ljungdahl pathway [64]. In contrast, the primary process in *heterotrophic* CO_2_ fixation is anaplerosis, the replenishment of tricarboxylic acid (TCA) cycle intermediates such as oxaloacetate and malate (see below). Autotrophic CO_2_ fixation pathways in prokaryotes (KEGG pathway ko00720) were enriched in both the nSM and vSM compared to no-DOM controls, but this was not statistically significant, and no clear difference was observed between the nSM and vSM (Figure S10). This suggests that chemoautotrophy is not the main process responsible for the observed DCF. Heterotrophic CO_2_ fixation, which is composed of various individual genes (see below), lacks a dedicated KEGG pathway, and therefore could not be assessed at the pathway level. However, the nSM metagenome was enriched in pathways related to carbohydrate utilization (e.g., the tricarboxylic acid (TCA) cycle and pyruvate metabolism), lipid metabolism (fatty acid metabolism) and amino acids synthesis (e.g., valine, leucine, and isoleucine biosynthesis) (Figure 3). We interpret these results as evidence that C-rich molecules such as carbohydrates and lipids were more abundant in the nSM compared with the vSM, consistent with the higher DOC:DON ratio.

**Figure 3.**
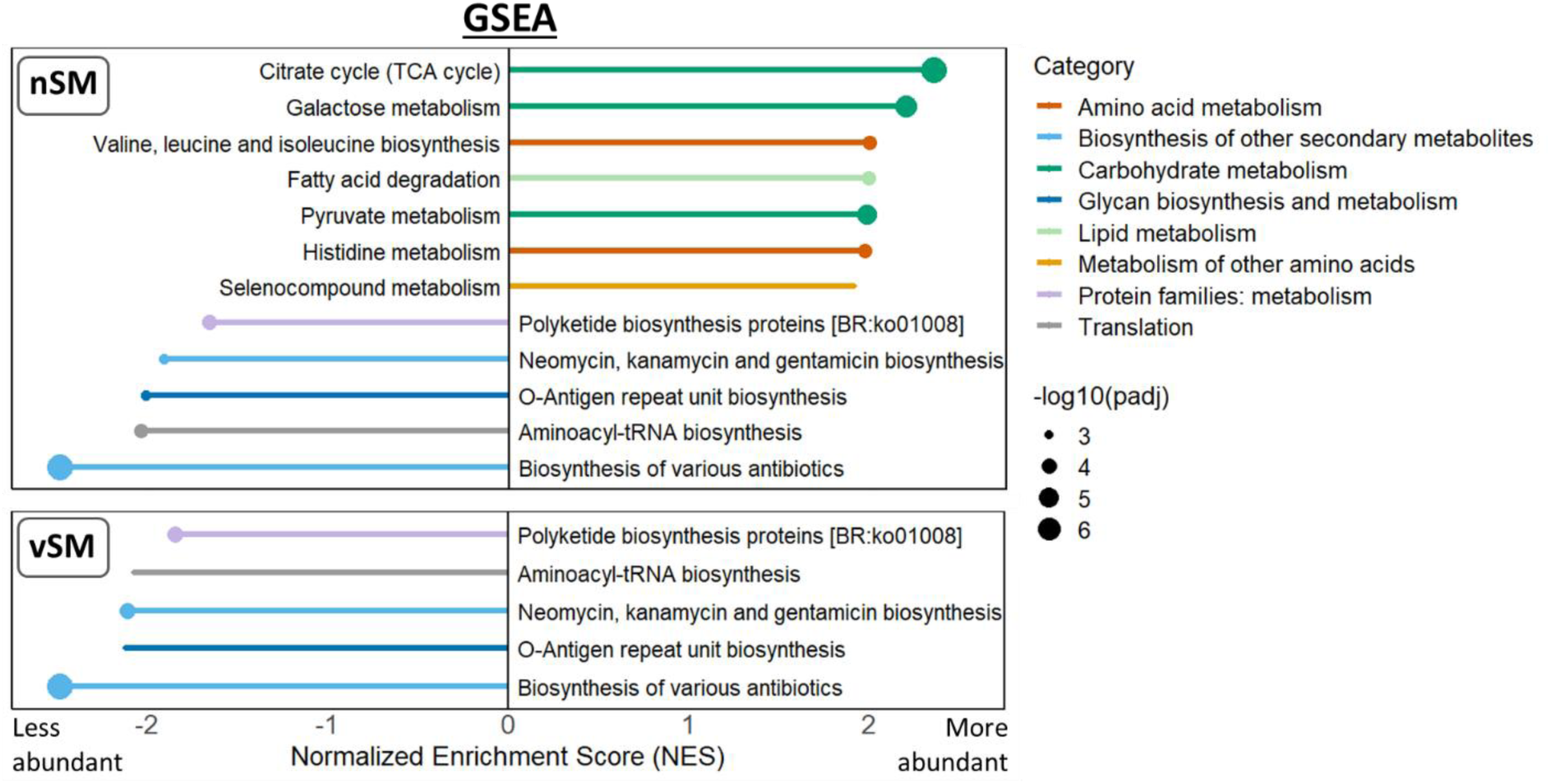
Gene set enrichment analysis (GSEA) of differentially expressed genes in nSM and vSM. Only statistically significant pathways (adjusted *p*-value < 0.05) are shown. The size of the bars represents the normalized enrichment score (NES). The size of the points represents the adjusted *p*-value. For expanded GSEA results see Figure S10.

If indeed carbon-rich molecules are major components of the nSM, organisms utilizing them as resources would likely need to produce their own nitrogen-rich building blocks such as amino acids and nucleotides from TCA cycle intermediates, rather than taking them up from the environment. In this case, the cell needs to compensate for flux diverted from the TCA cycle into biosynthesis by using one or more of six specific anaplerotic reactions, four of which also include a CO_2_ fixation step (Figure 4A, [60]). Genes encoding the CO_2_ fixing enzymes phosphoenolpyruvate (PEP) carboxylase (PEPC), PEP carboxykinase (PEPCK) and malate dehydrogenase (MDH) were relatively abundant in the metagenomes, whereas pyruvate carboxylase (PC) was less common (Figure 4B). Overall, there was no significant difference in the abundance of these genes between the nSM, vSM and controls (Figure 4B), yet in each treatment the sequences of these genes primarily originated from different bacterial orders (Figure 4C). For example, genes encoding PEPCK and MD from *Vibrionales* and *Rhodobacterales*, respectively, were enriched in the nSM (∼1.5-3.5 and ∼2-4.5-fold change compared to vSM), suggesting these clades might be involved in the observed DCF. In contrast, PEPC, PC and MD from *Pelagibacterales* were enriched in the vSM (∼1.4-2.8-fold change), as were PEPC from *Pseudoalteromonadales* (∼4-fold change), positioning these clades as less likely candidates for the observed DCF. Similar results were observed also for genes encoding enzymes in the glyoxylate shunt (malate synthase and isocitrate lyase), an anaplerotic process that does not include a CO_2_ fixation step (Figure 4A). Genes encoding other heterotrophic CO_2_ fixing enzymes, such as those involved in leucin utilization or biotin production, were less common in the metagenomes (Figure 4B).

**Figure 4.**
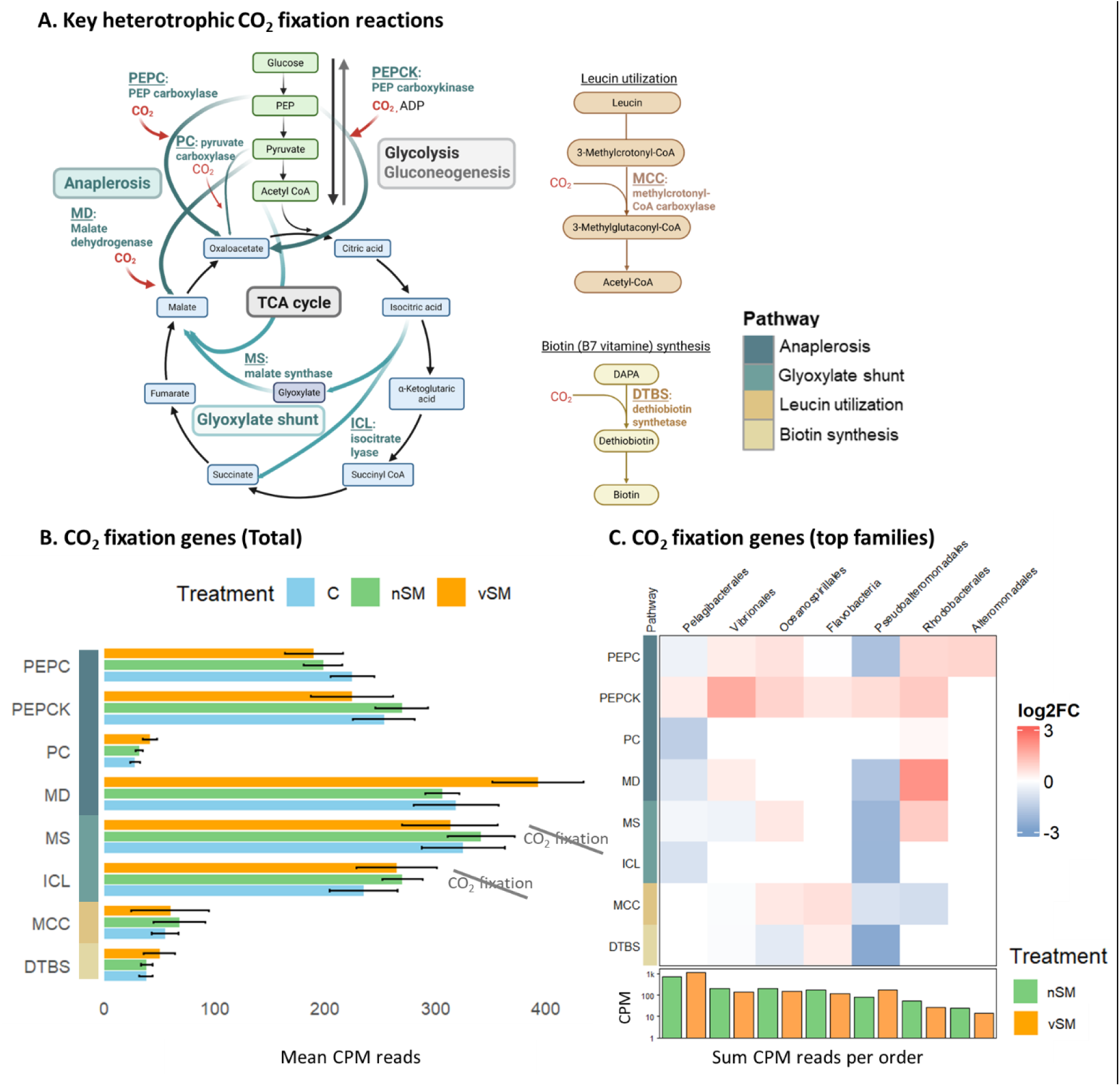
Metagenome analysis for CO2 fixation pathways. (A) Diagram illustrating key heterotrophic CO2 fixation reactions, along with the TCA, glyoxylate shunt and gluconeogenesis. DAPA: 7,8-diaminononanoate. Scheme was modified from Amano et al. [58]. (B) Total normalized reads (CPM)) for key enzymes associated with heterotrophic CO2 fixation. (C) Heatmap for average (N=3) fold change of nSM relatives to vSM reads for major heterotrophic taxa, which exhibited the highest relative abundance of discussing genes (based on CPM reads, Figure S12). Bottom bar plots showing sum CPM reads of all genes for each bacterial family in nSM (green) and vSM (orange). Abbreviations: PEPC - phosphoenolpyruvate (PEP) carboxylase; PEPCK - PEP carboxykinase; PC – pyruvate carboxylase; MDH - malate dehydrogenase; MS – malate synthase; ICL – isocitrate lyase; MCC - methylcrotonyl-CoA carboxylase; DTBS - dethiobiotin synthetase. Although ICL and MS have anaplerotic role, they are not involved in CO2 fixation – for more details see Supplementary text 3.

Analysis of metagenome-assembled genomes (MAGs) provided further support for hypothesis that heterotrophic, rather than autotrophic, processes were involved in the DCF, as pathways encoding the former were more commonly found and more complete within MAGs (Figure S11). Anaplerotic genes were present at varying levels of completeness across all MAGs, with *Rhodobacteraceae* MAG MED-G52 possessing all of the 4 CO_2_-fixing genes (Figure 5). However, the MAGs also further highlighted that there are other clades (primarily alpha and gamma proteobacteria and flavobacteria) that have the genetic potential to perform DCF.

**Figure 5.**
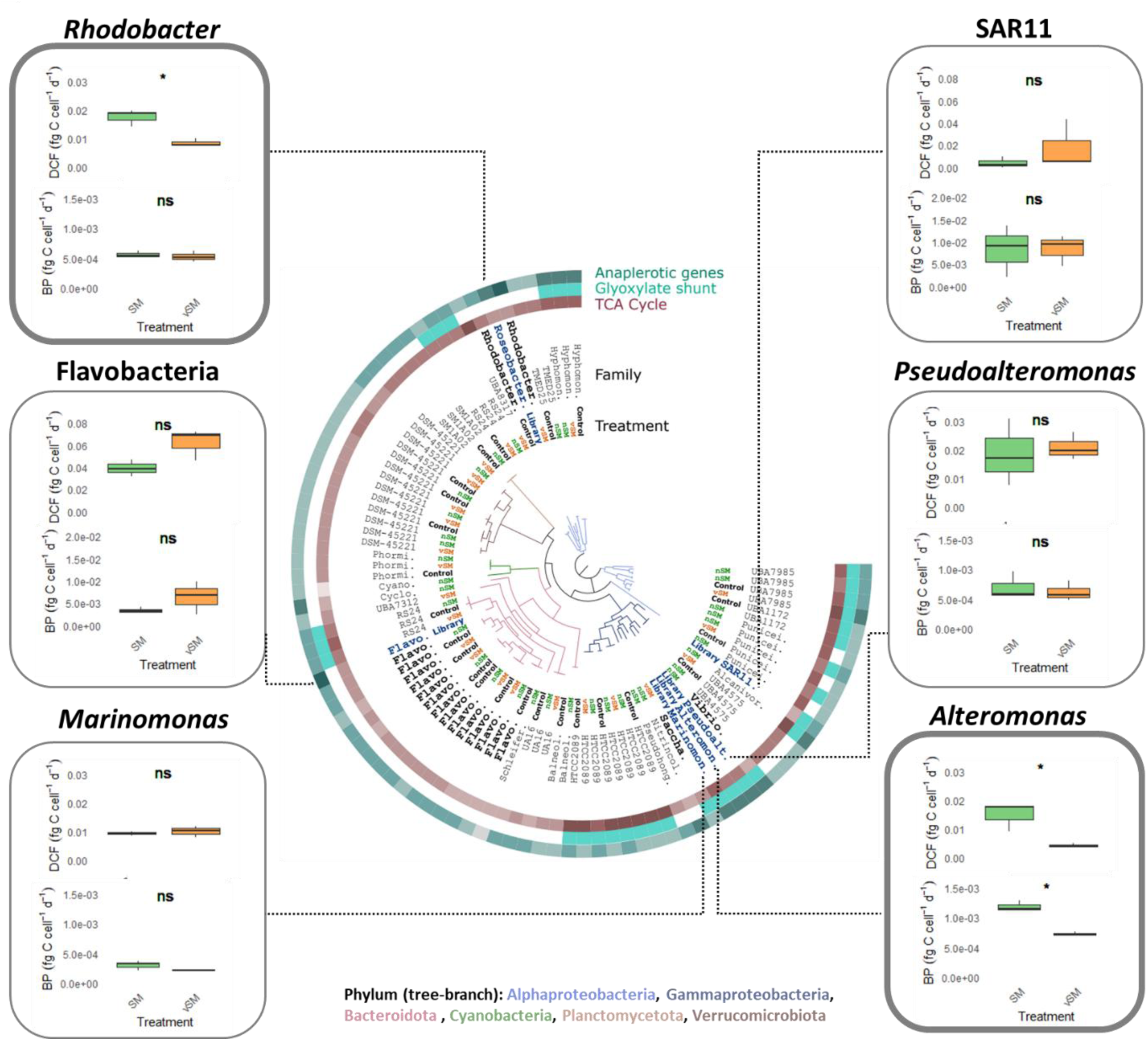
Phylogenomic tree illustrating metabolic potential of MAGs and activity in response to vSM and nSM of representative laboratory taxa. The outer rings indicate the presence of genes related to the TCA cycle, glyoxylate shunt, and anaplerotic pathways. Taxa are further annotated by bacterial family (inner ring) and treatment condition (control: black, nSM: green, vSM: orange). Bacterial taxa discussed in Figure 4C are highlighted in bold, and genomes derived from laboratory cultures are shown in blue. Boxplots adjacent to the tree illustrate per-cell dark inorganic carbon fixation (DCF) and bacterial production (BP) rates (in fg C cell⁻¹ d⁻¹) for selected laboratory strains that belong to the orders displayed in Figure 4C. An asterisk (*) denotes statistically significant results (*p*-value < 0.05), determined by a paired t-test with Bonferroni correction. Note the distinct Y-axis scales for *Flavobacteria* and SAR11 compared to other strains, likely attributable to their lower cell counts as measured by flow cytometry (Figure S13A). **Abbreviations for the family level:** Alcanivor. – *Alcanivoracaceae*; Alteromon. – *Alteromonadaceae*; Balneol. – *Balneolaceae*; Cyano. – *Cyanobiaceae*; Cyclo. – *Cyclobacteriaceae*; Flavo. – *Flavobacteriaceae*; Hyphomon. – *Hyphomonadaceae*; Marinomon. – *Marinomonadaceae*; Nitrincol. – *Nitrincolaceae*; Phormi. – *Phormidesmiaceae*; Pseudoalt. – *Pseudoalteromonadaceae*; Pseudohong. – *Pseudohongiellaceae*; Punicei. – *Puniceispirillaceae*; Rhodobacter. – *Rhodobacteraceae*; Saccha. – *Saccharospirillaceae*; Schleif. – *Schleiferiaceae*; Vibrio. – *Vibrionaceae*. **Abbreviations for the library strains (in blue):** Alteromon. – *Alteromonas mediterranea* DE; Flavo. – *Formosa agariphila* KMM; Marinomon. – *Marinomonas mediterranea* MMB1; Pseudoalt. – *Pseudoalteromonas haloplanktis* TAC; Roseobacter. – *Roseobacteraceae Marinovum* HOT5_F3 (order of *Rhodobacterales*); SAR11 - Candidatus pelagibacteraceae St. HTCC1062 (refer to Table S1 for full details).

### Cultured Rhodobacterales and Alteromonadales strains perform higher DCF in response to nSM compared to vSM

The metagenomic analysis highlights *Rhodobacterales* and *Alteromonadaceae* (as well as *Oceanospirillales* and *Vibrionales*) as orders potentially involved in the higher DCF in nSM treatment, and *Pseudoalteromonadales* and *Pelagibacterales* as relatively abundant clades less likely to do so (Figure 4C). To begin testing these hypotheses, six laboratory cultures, representing the major orders found in our metagenome analyses, were starved for carbon to simulate oligotrophic conditions followed by the addition of nSM and vSM (Supplementary text 4). Two of the strains, *Alteromonas mediterranea* strain DE (*Alteromonadales*) and *Marinovum* strain HOT5_F3 (*Rhodobacterales*), exhibited elevated per-cell DCF in the nSM (Figure 5), reminiscent of the observations in the field experiments (Figure 4C). None of the strain exhibited significantly higher DCF in response to vSM. Bacterial production also increased more in *Alteromonas* in response to nSM compared to vSM, and thus the DCF could, in principle, represent stronger overall growth. In contrast, only DCF increased in *Marinovum*, suggesting a specific metabolic response that is not coupled with growth, at least not in the time-frame of the experiment. This is similar to the response of the *Rhodobacterales* in the two experiments with natural microbial populations (Figure 3C).

### Summary and Conclusions

There are many ways microbial cells can die in the oceans. In laboratory cultures, whether *Prochlorococcus* MED4 dies due to phage infection or nitrogen starvation affects its biochemical structure, and the composition of DOM released upon death. In turn, these changes can have cascading effects on the microbes in the surrounding seawater, including shifts in bacterial community activity, metabolism, structure and function. DOM released through viral lysis increased overall heterotrophic community production, in agreement with a recent study [65], but we show for the first time that DOM from nutrient-starved cells increases heterotrophic CO_2_ fixation (Figure 6).

**Figure 6.**
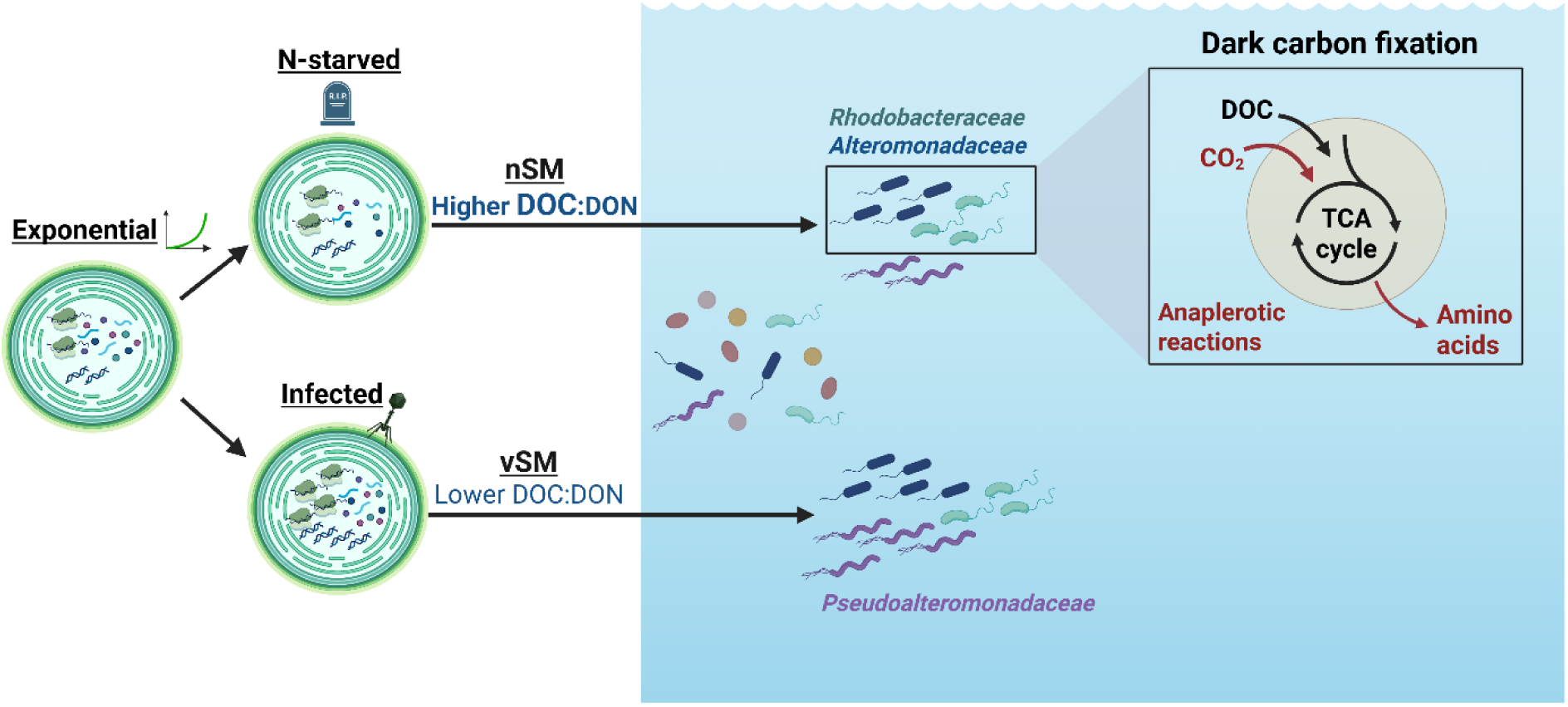
Summary illustration of experimental observations and suggested mechanism. Phage infection resulted in increased RNA and DNA quotas, whereas nitrogen-starved cells showed decreased protein quotas compared to logarithmic growth. Starved cells released more DOC concentration, while infected cells released more virions, found as dissolved DNA in the viral spent media (vSM). Addition of vSM resulted in increased total cells count and bacterial productivity. Addition of nSM increased dark inorganic carbon fixation (DCF) in both microcosm experiments and cultures of *Rhodobacterales and Alteromonadales*, which were also observed in the 16S rRNA analysis. Metagenomic analysis revealed an enrichment of pathways in the nSM treatments, including the TCA cycle, pyruvate metabolism, and phospholipid degradation.

In theory, phytoplankton biomass provides a “balanced meal” of building blocks (amino acids, nucleotides) and energy-rich molecules (carbohydrates, lipids) for heterotrophic bacteria [8,54]. However, the biochemical composition of the “meal” can change in response to different forms of death. Under nutrient starvation, phytoplankton like *Prochlorococcus* may shift towards producing and releasing excess carbon-rich metabolites, such as carbohydrates, through a process known as “overflow metabolism” [6,53,66]. Previous studies have shown that a significant portion of the carbohydrates released by *Prochlorococcus* are carboxylic acids such as pyruvate, glycolate, acetate and citrate (∼4-20%, [6,25–28,53,66]). These carboxylic acids either feed directly into the TCA cycle or are components of the cycle itself. We propose that such molecules (or their polymeric forms, e.g. lipids) comprise a larger part of the DOM released during starvation compared with phage lysis. We also propose that heterotrophic bacteria will need to employ anaplerotic reactions to enable them to utilize these carbon-rich molecules as resources for both energy and as backbones for biosynthesis of proteins and nucleic acids, e.g. using inorganic nitrogen sources. In support of this hypothesis, the DOC:DON ratio of the nSM was higher than that of the vSM, and pathways such as the TCA cycle, pyruvate metabolism, and fatty acid degradation were enriched in the community responding to the nSM (Figure 6).

Additionally, two of the clades shown to respond to nSM by growth and/or DCF, *Alteromonadales* and *Rhodobacterales,* respond in the lab and in the field to additions of organic acids, monosaccharides and disaccharides [30,67–70]. Amano et al. [58] also demonstrated that *Alteromonas* DE exhibits higher DCF rates on glucose compared to alanine, further supporting our hypothesis that carbon-rich DOM contributes to elevated DCF. Both of these clades are often associated with phytoplankton blooms and exudates, particularly those of cyanobacteria [71,72]. Taken together, these results, as well as previous studies, suggest an important role for these bacterial clades in anaplerotic CO_2_ fixation in the oceans [58,70,73,74].

In contrast to nitrogen starvation, cyanophage infection has been shown to increase the release of nitrogen-rich compounds like amino acids [1,22]. This suggests that DOM released by phage infection, and, potentially, phage themselves, are a source of bioavailable nitrogen, and potentially also other limiting nutrients such as phosphorus [75–77]. We propose that the larger increase in *Pseudoalteromonadaceae* abundance observed following vSM exposure in the autumn experiment was due to vSM’s nitrogen-rich composition, since cultured representatives of this clade demonstrated preferences for peptides in laboratory cultures and field incubations [30,67]. It is currently unclear whether the heterotrophic bacteria utilize nitrogen-rich lysate molecules such as cellular proteins or nucleic acids, or whether they take up and metabolize the phage themselves.

Our findings align with a global ocean metatranscriptomic study highlighted the ubiquitous expression of anaplerotic and glyoxylate shunt genes, often exceeding those involved in chemoautotrophy [58]. If, indeed, a fraction of this heterotrophic CO_2_ fixation is due to the presence of organic matter released due to nutrient starvation, this could have important consequences for understanding the cycling of dissolved organic carbon pools in the ocean. The relative contribution of different forms of death - grazing, phage infection and nutrient starvation – to the overall mortality of *Prochlorococcus* is still not well known [9–12]. However, it is intriguing that in the oligotrophic Gulf of Eilat, higher DCF rates occurred during the oligotrophic summer (∼12% vs. ∼7.5% in winter), supporting the link between nutrient scarcity and increased DCF [61]. It is also important to note that the increase in the released DOC:DON ratio does not necessarily require cell death, but can also occur as a result of overflow metabolism where fixed carbon is released when there is not enough N/P/Fe to fuel cell growth [78]. These processes (overflow metabolism and/or nutrient starvation) could be important processes responsible for increased DOC in the subtropical gyres [79]. Taken together, these results provide additional motivation to determine to what extent there exists a close coupling between specific forms of mortality, the release of DOM and its subsequent metabolism, the temporal succession of microbes in the sea (e.g. the succession of heterotrophic bacteria during phytoplankton blooms) and biogeochemical cycles.

## Materials and Methods

### *Prochlorococcus* bottle experiment: experimental setup and samples collection

Triplicate bottles containing 750 mL of *Prochlorococcus* MED4 were grown in low-nitrogen media (lowN, [43]): Mediterranean sweater (MSW) supplemented with 200µM NH_4_, 50µM PO_4_ and 10,000X trace metals [80]. After 5 days, half of the bottles were inoculated with cyanophage P-HM2 at MOI of 2:1 (viable phage particles were counted using plaque assay, see Supplementary text 5 for further details). P-HM2 is a T4-like myovirus with a latent period of approximately 5 hours [22,81]. Cells were infected on day 5, during early-to-mid logarithmic growth. Sampling occurred at three timepoints: T1 (∼12 hpi, following at least one round of infection), T2 (∼36 hpi, following several infection cycles), and T3 (uninfected samples only) captured the mid-decline stage for the N-starved culture (Figure 1B) (See Supplementary text 6 for more details on specific samples).

### Cruise and microcosm experimental set-up

Spent media from nitrogen-starved (nSM) and viral-infected (vSM) *Prochlorococcus* was introduced into surface seawater collected from the EMS. This experiment was conducted twice: once in autumn (November 7, 2022) and once in spring (March 28, 2024 during an IOLR transect cruise H51). Seawater for both experiments were collected at station H05 (32.99°N, 34.50°E), an open ocean site, from a depth of 25m. The experiment included 3 treatments: (1) “nSM”, amended with spent media from N-starved cells (T3 in the bottle experiment), (2) “vSM”, amended with lysate from infected cells (T2 in the bottle experiment) and a control amended only with inorganic N and P. Final DOC, NH_4_ and PO_4_ concentration was set to 25 µM, 15 µM and 10 µM, respectively - levels of N and P corresponding to the highest concentrations measured in the *Prochlorococcus*-derived DOM from either experimental replicate. Triplicate bottles of 4L (autumn) and 400mL (spring) from each treatment were incubated in dark for 24 hours.

### Measurements of bulk protein, RNA and DNA

Samples for protein, RNA and DNA were filtered on 0.7 μm GF/F filters and stored at −80℃ until analysis. Proteins were extracted by incubating the samples with lysis buffer at 37°C for 30 minutes, followed by 10 min of sonication in a water bath (Ultrasonic Cleaner AC-120H), and quantified using the bicinchoninic acid (BCA) assay [82]. RNA was isolated from the filter using Trizol extraction [83], and DNA using Phenol-Chloroform [84]. The concentration of both RNA and DNA was determined with the Qubit™ RNA HS Assay Kit and Qubit™ dsDNA HS Assay Kit, respectively.

### Measurements of NH_4_, PO_4_, DOC and DON in nSM and vSM

Samples for NH_4_ and PO_4_ were filtered through 0.2 μm polycarbonate filters and stored at −20℃ until analysis. Prior to measurements, samples were diluted 4-fold in DDW to reduce the salt level. NH_4_ was measured using an HI 96715 meter (Hanna instrument), and PO_4_ using Phosphate Colorimetric Assay Kit (Catalog Number MAK030). Samples for dissolved organic carbon (DOC) and total nitrogen (TN) were filtered through 0.2 μm polycarbonate filters, and are further elaborated on Supplementary text 7.

### Bacterial productivity and Dark carbon fixation

Bacterial productivity (BP) was estimated using the ³H-leucine incorporation method [85] as detailed by Reich et al. [86]. Dark inorganic carbon fixation (DCF) was measured using the ^14^C incorporation method [87], with few modifications [88], as described in details by Reich et al. [86].

### 16S RNA and DNA extraction, DNA digestion, and cDNA synthesis

The bacterial community structure and activity were assessed through 16S and 18S rRNA and rDNA sequencing. For each incubation bottle, 1–2 L of seawater was filtered onto 0.2 µm filters, preserved in lysis buffer, and stored at −80°C until extraction. RNA and DNA were extracted using the NucleoSpin RNA Stool and the PureLink™ Genomic DNA Mini Kit, respectively, according to the manufacturer’s instructions. RNA extraction included DNase treatment, followed by reverse transcription into cDNA using the iScript cDNA Synthesis Kit.

### 16S and 18S rRNA and rDNA sequencing

16S rRNA and rDNA sequencing was conducted using primers 515F and 926R [89], targeting ∼400 bp products. For 18S rRNA, primers Euk_1391F and EukBR were used, targeting ∼200 bp products. Sequencing protocols are described in detail in Givati et al. [30]. Briefly, amplicons were generated using a two-stage PCR protocol, and libraries were sequenced on an Illumina MiSeq platform (V3 chemistry, 2 × 300 bp reads) with a 15% phiX spike-in. Library preparation and sequencing were performed at the Genomics and Microbiome Core Facility, Rush University, IL, USA.

### 16S rRNA and rDNA sequence analysis

All sequences of both autumn and spring experiments were analyzed together using the Dada2 pipeline [90] and the software packages R [91] and Rstudio [92], as detailed in Givati et al. [30]. Forward and backward primers were trimmed, and forward and backward reads were truncated after quality inspections to 230 and 200 bases, respectively. After sequences merging, a consensus length of only between 353 and 366 bases was accepted, discarding ∼30% of sequences in the process. Finally, amplicon sequence variants (ASVs) that have less than 100 in total (all samples) were removed, remaining with a total of ∼720 ASVs. Silva database version 138 [93] was used for taxonomic assignment. All chloroplasts, mitochondria, archaea, eukaryotes, and ASVs without any taxonomic affiliation were discarded from downstream analyses. Full ASVs and taxonomy list of 16S rRNA and rDNA can be found in Supplementary files 1 and 2. For details on 18S rRNA sequence analysis see Supplementary text 8.

### Metagenomics sequencing

Shotgun metagenomic libraries were prepared using the Illumina Nextera XT DNA Library Preparation Kit. A total of 9 samples were processed for dsDNA shotgun library preparation. Sequencing was carried out on an Illumina NovaSeq X platform using a 2 × 150 bp paired-end protocol. Approximately 900 million clusters were generated per lane on a 25B flow cell.

### Metagenomics and MAGs construction

Raw metagenomic reads were first assessed for quality using FastQC [94], followed by quality filtering and trimming using the algorithm bbduk [95]. For *De-novo* assembly, trimmed metagenomes were assembled using Megahit [96]. For each biological treatment (Control/SM/vSM), we first pooled the replicate libraries and then performed a combined assembly. Gene calls on each of the assembly files were made using [97]. All the gene-level KEGG Ortholog (KO) functional annotation was performed using KOFamScan [98]. Gene- and contig-level taxonomic classifications for each assembly were performed using CAT [99]. Trimmed short reads for each sample were mapped to the corresponding assemblies (contigs/genes) and coverage information was generated using bowtie2 [100]. All gene- and taxonomy-level CPM read counts are provided in Supplementary files 4 and 5 (combined) and in Supplementary files 6 to 8 (separated by treatment). MAGs were reconstructed by binning assembled contigs per condition (see Supplementary text 9 for more details).

### Differential Gene Set Enrichment Analysis (GSEA)

Differential Gene Set Enrichment Analysis (GSEA) was performed using the fgsea R package [101]. Gene sets were defined based on the KEGG pathway database, utilizing a table containing KO IDs, their corresponding functions, and pathway assignments. Analysis was restricted to KO IDs identified in the metagenome and metabolic pathways containing 10 to 500 KOs. GSEA was conducted with 10,000 permutations, using fold-change values as the ranking metric. Gene sets with an adjusted *p*-value < 0.05 were considered statistically significant.

### Statistics

Data pre-processing and statistical analyses were performed using the R statistical programming language [91,92] and on Linux ubunto20. One-way analysis of variance (ANOVA) and post hoc Tukey test was performed to compare the treatments in lab and microcosms experiments (Figure 2 and Figure 3) using multcompView [102] and dplyr [103] packages. Paired t-test with Bonferroni correction was used to compare the means between treatments for each bacteria (Figure 6) using the packages tidyverse v1.3.0 [104], rstatix [105], and ggpubr [106]. Microbial community differences between treatments in the NMDS plots were statistically assessed by PERMANOVA on Bray-Curtis dissimilarity matrices, with 999 permutations to determine significance using vegan package [107]. Plots were originated using the ggplot2 package [108].

## Data availability

All forward and backward sequence reads were uploaded to NCBI under umbrella project accession number PRJNA1291108: PRJNA1287306 (16S rRNA), PRJNA1287330 (16S rDNA), PRJNA1291108 (18S rRNA) and PRJNA1302148 (metagenomics).

## Acknowledgements

We thank Ximena Dubinsky for help with the BP measurements, Solvig Pinnow and Hans-Peter Grossart for the DON measurements, Eyal Giesler and Edo Bar Zeev for the DOC measurements, Anat Tsemel and Mike Krom for the inorganic nutrient measurements. We also thank Abdiel Lázaro García and Laura Steindler for the SAR11 cultures, and Ankur Naqib, Adit Chaudhary and Stefan Green from the Rush Research Bioinformatics Core for the metagenomic analysis pipeline. This study was supported by the Israel Science Foundation (grant number 1786/20 to DS) and by the National Science Foundation - United States-Israel Binational Science Foundation (NSFOCE-BSF 1635070 and NSF-BSF 2246707 to DS).

## References

1. Cai L, Li H, Deng J et al. Biological interactions with Prochlorococcus: implications for the marine carbon cycle. Trends Microbiol 2024;32:280–91.

2. Biller SJ, Lundeen RA, Hmelo LR et al. Prochlorococcus extracellular vesicles: molecular composition and adsorption to diverse microbes. Environ Microbiol 2022;24:420–35.

3. Biller SJ, Schubotz F, Roggensack SE et al. Bacterial Vesicles in Marine Ecosystems. Science (80-) 2014;343:183–6.

4. Biller SJ, Berube PM, Lindell D et al. Prochlorococcus: the structure and function of collective diversity. Nat Rev Microbiol 2015;13:13–27.

5. Flombaum P, Gallegos JL, Gordillo RA et al. Present and future global distributions of the marine Cyanobacteria Prochlorococcus and Synechococcus. Proc Natl Acad Sci 2013;110:9824–9.

6. Bertilsson S, Berglund O, Pullin MJ et al. Release of dissolved organic matter by Prochlorococcus. Vie Milieu/Life Environ 2005:225–31.

7. Roth-Rosenberg D, Aharonovich D, Omta AW et al. Dynamic macromolecular composition and high exudation rates in Prochlorococcus. Limnol Oceanogr 2021;66:1759–73.

8. Thornton DCO. Dissolved organic matter (DOM) release by phytoplankton in the contemporary and future ocean. Eur J Phycol 2014;49:20–46.

9. Matteson AR, Rowe JM, Ponsero AJ et al. High abundances of cyanomyoviruses in marine ecosystems demonstrate ecological relevance. FEMS Microbiol Ecol 2013;84:223–34.

10. Carlson MCG, Ribalet F, Maidanik I et al. Viruses affect picocyanobacterial abundance and biogeography in the North Pacific Ocean. Nat Microbiol 2022;7:570–80.

11. Mruwat N, Carlson MCG, Goldin S et al. A single-cell polony method reveals low levels of infected Prochlorococcus in oligotrophic waters despite high cyanophage abundances. ISME J 2021;15:41–54.

12. Beckett SJ, Demory D, Coenen AR et al. Disentangling top-down drivers of mortality underlying diel population dynamics of Prochlorococcus in the North Pacific Subtropical Gyre. Nat Commun 2024;15:2105.

13. Ribalet F, Swalwell J, Clayton S et al. Light-driven synchrony of Prochlorococcus growth and mortality in the subtropical Pacific gyre. Proc Natl Acad Sci 2015;112:7–11.

14. Roth-rosenberg D, Aharonovich D, Omta A et al. Dynamic macromolecular composition and high exudation rates in Prochlorococcus. bioRxiv 2020.

15. Casey JR, Boiteau RM, Engqvist MKM et al. Basin-scale biogeography of marine phytoplankton reflects cellular-scale optimization of metabolism and physiology. Sci Adv 2022;8:eabl4930.

16. Geider RJ, La Roche J. Redfield revisited: Variability of C:N:P in marine microalgae and its biochemical basis. Eur J Phycol 2002;37:1–17.

17. Liefer JD, Garg A, Fyfe MH et al. The macromolecular basis of phytoplankton C:N:P under nitrogen starvation. Front Microbiol 2019;10:1–16.

18. Jover LF, Effler TC, Buchan A et al. The elemental composition of virus particles: Implications for marine biogeochemical cycles. Nat Rev Microbiol 2014;12:519–28.

19. Thompson LR, Zeng Q, Kelly L et al. Phage auxiliary metabolic genes and the redirection of cyanobacterial host carbon metabolism. Proc Natl Acad Sci U S A 2011;108:E757–64.

20. Ankrah NYD, May AL, Middleton JL et al. Phage infection of an environmentally relevant marine bacterium alters host metabolism and lysate composition. ISME J 2014;8:1089–100.

21. Chevallereau A, Blasdel BG, De Smet J et al. Next-Generation “-omics” Approaches Reveal a Massive Alteration of Host RNA Metabolism during Bacteriophage Infection of Pseudomonas aeruginosa. PLoS Genet 2016;12:1–20.

22. Xiao X, Guo W, Li X et al. Viral Lysis Alters the Optical Properties and Biological Availability of Dissolved Organic Matter Derived from Prochlorococcus Picocyanobacteria. Johnson KN (ed.). Appl Environ Microbiol 2021;87:1–19.

23. Fang X, Liu Y, Zhao Y et al. Transcriptomic responses of the marine cyanobacterium Prochlorococcus to viral lysis products. Environ Microbiol 2019;21:2015–28.

24. Azam F, Malfatti F. Microbial structuring of marine ecosystems. Nat Rev Microbiol 2007;5:782–91.

25. Biller SJ, Coe A, Roggensack SE et al. Heterotroph Interactions Alter Prochlorococcus Transcriptome Dynamics during Extended Periods of Darkness. Mason O (ed.). mSystems 2018;3:1–18.

26. Braakman R, Follows MJ, Chisholm SW. Metabolic evolution and the self-organization of ecosystems. Proc Natl Acad Sci 2017;114:E3091–100.

27. Calfee BC, Glasgo LD, Zinser ER. Prochlorococcus Exudate Stimulates Heterotrophic Bacterial Competition with Rival Phytoplankton for Available Nitrogen. Martiny JBH (ed.). MBio 2022;13, DOI: 10.1128/mbio.02571-21.

28. Becker JW, Hogle SL, Rosendo K et al. Co-culture and biogeography of Prochlorococcus and SAR11. ISME J 2019;13:1506–19.

29. Kujawinski EB, Braakman R, Longnecker K et al. Metabolite diversity among representatives of divergent Prochlorococcus ecotypes. Gilbert JA (ed.). mSystems 2023;8, DOI: 10.1128/msystems.01261-22.

30. Givati S, Forchielli E, Aharonovich D et al. Diversity in the utilization of different molecular classes of dissolved organic matter by heterotrophic marine bacteria. Biddle JF (ed.). Appl Environ Microbiol 2024;90:e00256–24.

31. Haider MN, Iqbal MM, Nishimura M et al. Bacterial response to glucose addition: growth and community structure in seawater microcosms from North Pacific Ocean. Sci Rep 2023;13:341.

32. Eilers H, Pernthaler J, Amann R. Succession of Pelagic Marine Bacteria during Enrichment: a Close Look at Cultivation-Induced Shifts. Appl Environ Microbiol 2000;66:4634–40.

33. Baltar F, Lundin D, Palovaara J et al. Prokaryotic Responses to Ammonium and Organic Carbon Reveal Alternative CO2 Fixation Pathways and Importance of Alkaline Phosphatase in the Mesopelagic North Atlantic. Front Microbiol 2016;7:1–19.

34. Weitz JS, Stock CA, Wilhelm SW et al. A multitrophic model to quantify the effects of marine viruses on microbial food webs and ecosystem processes. ISME J 2015;9:1352–64.

35. Ma X, Coleman ML, Waldbauer JR. Distinct molecular signatures in dissolved organic matter produced by viral lysis of marine cyanobacteria. Environ Microbiol 2018;20:3001–11.

36. Weinbauer MG, Bonilla-findji O, Chan AMYM et al. Synechococcus growth in the ocean may depend on the lysis of heterotrophic bacteria. J Plankton Res 2011;33:1465–76.

37. Pourtois J, Tarnita CE, Bonachela JA. Impact of Lytic Phages on Phosphorus- vs. Nitrogen-Limited Marine Microbes. Front Microbiol 2020;11, DOI: 10.3389/fmicb.2020.00221.

38. Weinbauer MG. Ecology of prokaryotic viruses. FEMS Microbiol Rev 2004;28:127–81.

39. Shelford EJ, Middelboe M, Møller EF et al. Virus-driven nitrogen cycling enhances phytoplankton growth. Aquat Microb Ecol 2012;66:41–6.

40. Sterner R., Elser J. Ecological Stoichiometry: The Biology of Elements from Molecules to the Biosphere., 2002.

41. Moreno AR, Martiny AC. Ecological Stoichiometry of Ocean Plankton. Ann Rev Mar Sci 2018;10:43–69.

42. Sher D, Segrè D, Follows MJ. Quantitative principles of microbial metabolism shared across scales. Nat Microbiol 2024;9:1940–53.

43. Grossowicz M, Roth-rosenberg D, Aharonovich D et al. Prochlorococcus in the lab and in silico : The importance of representing exudation. 2017, DOI: 10.1002/lno.10463.

44. Ni T, Zeng Q. Diel Infection of Cyanobacteria by Cyanophages. Front Mar Sci 2016;2:1–7.

45. Blazewicz SJ, Barnard RL, Daly RA et al. Evaluating rRNA as an indicator of microbial activity in environmental communities: Limitations and uses. ISME J 2013;7:2061–8.

46. Breitbart M, Thompson LR, Suttle CA et al. Exploring the vast diversity of marine viruses. Oceanography 2007;20:135–9.

47. Thompson LR, Zeng Q, Chisholm SW. Gene Expression Patterns during Light and Dark Infection of Prochlorococcus by Cyanophage. Campbell DA (ed.). PLoS One 2016;11:e0165375.

48. Mizuno CM, Guyomar C, Roux S et al. Numerous cultivated and uncultivated viruses encode ribosomal proteins. Nat Commun 2019;10:752.

49. Chen L-X, Jaffe AL, Borges AL et al. Phage-encoded ribosomal protein S21 expression is linked to late-stage phage replication. ISME Commun 2022;2:1–10.

50. Brüwer JD, Sidhu C, Zhao Y et al. Globally occurring pelagiphage infections create ribosome-deprived cells. Nat Commun 2024;15:3715.

51. Laurenceau R, Raho N, Forget M et al. Frequency of mispackaging of Prochlorococcus DNA by cyanophage. ISME J 2021;15:129–40.

52. Moran MA, Kujawinski EB, Stubbins A et al. Deciphering ocean carbon in a changing world. Proc Natl Acad Sci U S A 2016;113:3143–51.

53. Szul MJ, Dearth SP, Campagna SR et al. Carbon Fate and Flux in Prochlorococcus under Nitrogen Limitation. Gutierrez M (ed.). mSystems 2019;4:1–16.

54. Eigemann F, Rahav E, Grossart H et al. Phytoplankton exudates provide full nutrition to a subset of accompanying heterotrophic bacteria via carbon, nitrogen and phosphorus allocation. Environ Microbiol 2022;24:2467–83.

55. Hu W, Zhang H, Lin X et al. Characteristics, Biodiversity, and Cultivation Strategy of Low Nucleic Acid Content Bacteria. Front Microbiol 2022;13, DOI: 10.3389/fmicb.2022.900669.

56. Lebaron P, Servais P, Agogue H et al. Does the High Nucleic Acid Content of Individual Bacterial Cells Allow Us To Discriminate between Active Cells and Inactive Cells in Aquatic Systems ? Appl Environ Microbiol 2001;67:1775–82.

57. Alothman A, López-Sandoval D, Duarte CM et al. Bacterioplankton dark CO 2 fixation in oligotrophic waters. Biogeosciences 2023;20:3613–24.

58. Amano C, Willhelm U, Reinthaler T et al. Anaplerotic processes are key contributors to dark carbon xation in the ocean. 2024.

59. Baltar F, Herndl GJ. Ideas and perspectives: Is dark carbon fixation relevant for oceanic primary production estimates? Biogeosciences 2019;16:3793–9.

60. Braun A, Spona-Friedl M, Avramov M et al. Reviews and syntheses: Heterotrophic fixation of inorganic carbon – significant but invisible flux in environmental carbon cycling. Biogeosciences 2021;18:3689–700.

61. Reich T, Belkin N, Sisma-Ventura G et al. Significant dark inorganic carbon fixation in the euphotic zone of an oligotrophic sea. Limnol Oceanogr 2024;69:1129–42.

62. Spona-Friedl M, Braun A, Huber C et al. Substrate-dependent CO2 fixation in heterotrophic bacteria revealed by stable isotope labelling. FEMS Microbiol Ecol 2020;96:1–13.

63. Eigemann F, Rahav E, Grossart HP et al. Phytoplankton Producer Species and Transformation of Released Compounds over Time Define Bacterial Communities following Phytoplankton Dissolved Organic Matter Pulses. Appl Environ Microbiol 2023;89, DOI: 10.1128/aem.00539-23.

64. Hügler, M., & Sievert SM. Beyond the Calvin Cycle: Autotrophic Carbon Fixation in the Ocean. Ann Rev Mar Sci 2011;3:261–89.

65. Kranzler CF, Busono DA, Walsh GJ et al. Taxonomically distinct diatom viruses differentially impact microbial processing of organic matter. Sci Adv 2025;11, DOI: 10.1126/sciadv.adq5439.

66. Ofaim S, Sulheim S, Almaas E et al. Dynamic Allocation of Carbon Storage and Nutrient-Dependent Exudation in a Revised Genome-Scale Model of Prochlorococcus. Front Genet 2021;12:1–21.

67. Forchielli, E., Sher, D., & Segrè D. Metabolic phenotyping of marine heterotrophs on refactored media reveals diverse metabolic adaptations and lifestyle strategies. Msystems 2022;7:e00070–22.

68. Landa M, Burns AS, Roth SJ et al. Bacterial transcriptome remodeling during sequential co-culture with a marine dinoflagellate and diatom. ISME J 2017;11:2677–90.

69. Moreno-Cabezuelo JÁ, Gómez-Baena G, Díez J et al. Integrated Proteomic and Metabolomic Analyses Show Differential Effects of Glucose Availability in Marine Synechococcus and Prochlorococcus. Hom EFY (ed.). Microbiol Spectr 2023;11, DOI: 10.1128/spectrum.03275-22.

70. Swingley WD, Sadekar S, Mastrian SD et al. The Complete Genome Sequence of Roseobacter denitrificans Reveals a Mixotrophic Rather than Photosynthetic Metabolism. J Bacteriol 2007;189:683–90.

71. Ferrer-González FX, Widner B, Holderman NR et al. Resource partitioning of phytoplankton metabolites that support bacterial heterotrophy. ISME J 2021;15:762–73.

72. Kieft B, Li Z, Bryson S et al. Phytoplankton exudates and lysates support distinct microbial consortia with specialized metabolic and ecophysiological traits. Proc Natl Acad Sci 2021;118:1–12.

73. Moran MA, Belas R, Schell MA et al. Ecological Genomics of Marine Roseobacters. Appl Environ Microbiol 2007;73:4559–69.

74. Hameed A, Suchithra KV, Lin S-Y et al. Genomic potential for inorganic carbon sequestration and xenobiotic degradation in marine bacterium Youngimonas vesicularis CC-AMW-ET affiliated to family Paracoccaceae. Antonie Van Leeuwenhoek 2023;116:1247–59.

75. Noble R, Fuhrman J. Breakdown and microbial uptake of marine viruses and other lysis products. Aquat Microb Ecol 1999;20:1–11.

76. Mayers KMJ, Kuhlisch C, Basso JTR et al. Grazing on Marine Viruses and Its Biogeochemical Implications. Prasad VR (ed.). MBio 2023;14, DOI: 10.1128/mbio.01921-21.

77. Middelboe M, Jørgensen NOG. Viral lysis of bacteria: an important source of dissolved amino acids and cell wall compounds. J Mar Biol Assoc United Kingdom 2006;86:605–12.

78. Dubinsky Z, Berman-Frank I. Uncoupling primary production from population growth in photosynthesizing organisms in aquatic ecosystems. Aquat Sci 2001;63:4–17.

79. Wu Z, Dutkiewicz S, Jahn O et al. Modeling Photosynthesis and Exudation in Subtropical Oceans. Global Biogeochem Cycles 2021;35, DOI: 10.1029/2021GB006941.

80. Moore LR, Coe A, Zinser ER et al. Culturing the marine cyanobacterium Prochlorococcus. Limnol Oceanogr Methods 2007;5:353–62.

81. Sullivan MB, Huang KH, Ignacio-Espinoza JC et al. Genomic analysis of oceanic cyanobacterial myoviruses compared with T4-like myoviruses from diverse hosts and environments. Environ Microbiol 2010;12:3035–56.

82. Smith PK, Krohn RI, Hermanson GT et al. Measurement of protein using bicinchoninic acid. Anal Biochem 1985;150:76–85.

83. Chomczynski P, Sacchi N. Single-step method of RNA isolation by acid guanidinium thiocyanate-phenol-chloroform extraction. Anal Biochem 1987;1:581–5.

84. Marmur J. A procedure for the isolation of deoxyribonucleic acid from micro-organisms. J Mol Biol 1961;3:208–IN1.

85. Simon M, Azam F. Protein content and protein synthesis rates of planktonic marine bacteria. Mar Ecol Prog Ser 1989;51:201–13.

86. Reich T, Ben-Ezra T, Belkin N et al. A year in the life of the Eastern Mediterranean: Monthly dynamics of phytoplankton and bacterioplankton in an ultra-oligotrophic sea. Deep Sea Res Part I Oceanogr Res Pap 2022;182:103720.

87. Nielsen ES. The Use of Radio-active Carbon (C14) for Measuring Organic Production in the Sea. ICES J Mar Sci 1952;18:117–40.

88. Hazan O, Silverman J, Sisma-Ventura G et al. Mesopelagic Prokaryotes Alter Surface Phytoplankton Production during Simulated Deep Mixing Experiments in Eastern Mediterranean Sea Waters. Front Mar Sci 2018;5, DOI: 10.3389/fmars.2018.00001.

89. Walters W, Hyde ER, Berg-lyons D et al. Transcribed Spacer Marker Gene Primers for Microbial Community Surveys. mSystems 2015;1:e0009–15.

90. Callahan BJ, McMurdie PJ, Rosen MJ et al. DADA2: High-resolution sample inference from Illumina amplicon data. Nat Methods 2016;13:581–3.

91. R Development Core Team RFFSC. R: A language and environment for statistical computing. 2011.

92. Team R. RStudio: integrated development for R. RStudio, PBC. 2020.

93. Quast C, Pruesse E, Yilmaz P et al. The SILVA ribosomal RNA gene database project: improved data processing and web-based tools. Nucleic Acids Res 2012;41:D590–6.

94. Andrews S. FastQC: a quality control tool for high throughput sequence data. 2010.

95. Bushnell B, Rood J, Singer E. BBMerge – Accurate paired shotgun read merging via overlap. Biggs PJ (ed.). PLoS One 2017;12:e0185056.

96. Li D, Liu C-M, Luo R et al. MEGAHIT: an ultra-fast single-node solution for large and complex metagenomics assembly via succinct de Bruijn graph. Bioinformatics 2015;31:1674–6.

97. Hyatt D, Chen G-L, LoCascio PF et al. Prodigal: prokaryotic gene recognition and translation initiation site identification. BMC Bioinformatics 2010;11:119.

98. Aramaki T, Blanc-Mathieu R, Endo H et al. KofamKOALA: KEGG Ortholog assignment based on profile HMM and adaptive score threshold. Valencia A (ed.). Bioinformatics 2020;36:2251–2.

99. von Meijenfeldt FAB, Arkhipova K, Cambuy DD et al. Robust taxonomic classification of uncharted microbial sequences and bins with CAT and BAT. Genome Biol 2019;20:217.

100. Langmead B, Salzberg SL. Fast gapped-read alignment with Bowtie 2. Nat Methods 2012;9:357–9.

101. Korotkevich G, Sukhov V, Budin N et al. Fast gene set enrichment analysis. bioRxiv 2016, DOI: 10.1101/060012.

102. Spencer Graves, Hans-Peter Piepho LS with help from SD-R. multcompView: Visualizations of Paired Comparisons. 2019.

103. Hadley Wickham, Romain François, Lionel Henry KM and DV. dplyr: A Grammar of Data Manipulation. 2023.

104. Wickham, H., Averick, M., Bryan, J., Chang, W., McGowan, L., François, R., Grolemund, G., Hayes, A., Henry, L., Hester, J., Kuhn, M., Pedersen, T., Miller, E., Bache, S., Müller, K., Ooms, J., Robinson, D., Seidel, D., Spinu, V., … Yutani, H. Wickham, H., H. Welcome to the tidyverse. J Open Source Softw 2019;4:1686.

105. Kassambara A. rstatix: Pipe-Friendly Framework for Basic Statistical Tests. 2023.

106. Kassambara A. ggpubr: “ggplot2” Based Publication Ready Plots. 2023.

107. Jari Oksanen, Gavin L. Simpson, F. Guillaume Blanchet, Roeland Kindt P, Legendre, Peter R. Minchin, R.B. O’Hara, Peter Solymos MHHS, Eduard Szoecs, Helene Wagner, Matt Barbour, Michael Bedward BB et al. vegan: Community Ecology Package. 2022.

108. Wickham H. ggplot2: Elegant Graphics for Data Analysis. 2016.

