## Supplementary text and figures for "Distinct microbial response to organic matter from nitrogen-starved and virus-infected phytoplankton"

### Supplementary Information

#### Supplementary text 1 – comparing DCF with primary production rates

DCF rates in the nSM treatment (~20 µg C $L^{-1}$ $d^{-1}$) increased approximately 75-and 100-fold relative to $T_0$ (~0.25 µg C $L^{-1}$ $d^{-1}$ in the spring and ~0.1 µg C $L^{-1}$ $d^{-1}$ in the autumn, respectively), while total cells increased by only ~2-3-fold. The low $T_0$ values align with previous measurements of primary productivity rates (photosynthesis) in EMS, which range from ~0 to 3.5 µg C $L^{-1}$ $d^{-1}$ [1]. In contrast, the elevated DCF correspond to highest primary productivity measured in the surface water from the Gulf of Aqaba [2]. These rates fall within the range reported in other marine environments, including more eutrophic regions such as tropical estuaries [3,4]. The substantial values of DCF measured here support the significant role of DCF in the ocean, highlighting the contribution of heterotrophic bacteria to total $CO_2$ fixation. It should be noted that DCF also increased in the control amended only with inorganic nutrients. We speculate that the addition of N and P to the already starved population allowed them to utilize existing carbon sources, thus prompting DCF.

**Supplementary text 2 – comparing microbial community structure between two experiments**

The initial microbial community composition differed significantly between autumn and spring, as anticipated (Figure S6). Oligotrophic bacteria, like members of AEGEAN-169, SAR11 clade 1 and SAR116 were more prevalent in the autumn. Their relative abundance declined with nutrient addition and decreased even further with the introduction of organic. Contrary, the relative abundance of other Alphaproteobacteria, like *Hyphomonadaceae* and *Rhodobacteraceae*, *Flavobacteriaceae* and members of the Gammaproteobacteria increased along nutrients and organic matter addition. Overall, the differences between the treatments, control, and $T_0$ were more pronounced in spring, where the control diverged from $T_0$, and the nSM and vSM treatments clustered together (Figure S6A).

As opposed to our initial hypothesis, no major differences were observed in the resulting community structure between the treatments in both experiments. *Rhodobacteraceae* doubled their relative abundance in both experiments in response to nSM and vSM, from approximately 3% to 6% and from ~6 to ~12% compared to the control (Figure S6B). In both seasons, a single dominant ASV was present, accounting for approximately 3% and 6% of total reads in nSM and vSM, respectively. Although these ASVs were different between seasons, each shared 94.46% identity based on BLAST (NCBI) and could not be identified to the genus level. They were also detected in the control but at lower abundances (1.5% and 0.5%, respectively). Another group that significantly increased in both treatments was *Ateromonadaceae* in the spring, reaching a relative abundance of approximately 14% in the nSM treatment and 20% in the vSM compared to ~4.5% in the control (Figure S6B). The *Ateromonadaceae* consisted of 3-4 mostly abundant ASVs, with no difference between the treatments. Furthermore, *Pseudoalteromonadaceae* primarily responded to vSM in the autumn, with an average relative abundance of ~7% (Figure S6B). One ASV was predominantly within *Pseudoalteromonadaceae*, accounting for ~5.5% of reads in the vSM samples. In spring, the same ASV was also the most dominant in both treatments (~1%), despite no significant growth of *Pseudoalteromonadaceae* being observed.

Were these bacteria abundant in the original seawater? The Alphaproteobacteria *Rhodobacteraceae* was already identified in $T_0$ (~3-4% in both seasons), while the Gammaproteobacteria *Ateromonadaceae* and *Pseudoalteromonadaceae* composed only ~0.1-0.2% from the total microbial community. Such Gammaproteobacteria species tends to bloom in diverse microcosms experiments, for example in response to specific organic molecules [5–8], phytoplankton SM [9–12] and phytoplankton vSM [13,14]. Moreover, they are widely associated with phytoplankton blooms [15,16].

Compared with the 16S rRNA, the changes in 16S rDNA were more subtle (Figure S8). Our investigation into the eukaryotic population (using 18S rRNA gene sequencing, Figure S9) indicated that the community structure remained consistent across all experimental treatments, the control, and the original seawater sample. To note, it is possible that the lack of significant changes in community structure was influenced by the relatively short 24-hour incubation period. A comparable study on *Prochlorococcus* MED4 SM and vSM found that most changes in community structure occurred within 36 hours [13].

**Supplementary text 3– glyoxylate shunt genes**

Two additional anaplerotic genes are isocitrate lyase (ICL) and malate synthase (MS). Although they play an anaplerotic role in the TCA cycle, they are not involved in $CO_2$ fixation [17]. Similar to other anplerotic genes, they exhibited high metagenomic coverage (Figure 4B). They were enriched in *Vibrionales*, *Oceanospirillales*, and *Pseudoalteromonadales* (Figure S10).

Additionally, the glyoxylate shunt pathway was identified in both Alphaproteobacteria and Gammaproteobacteria MAGs (e.g., most *Pseudomonadales* and *Vibrionaceae*), with all possessing a complete pathway (Figure 5).

**Supplementary text 4 - Lab-cultures experiment with heterotopic bacteria**

Bacterial cultures were revived in 2 mL of marine broth (MB) at 26°C for 48 hours in 12 mL sterile growth tubes, followed by several transfers to induced carbon starvation: 500µl of cultures were then transferred to 5mL ProMM medium and incubated for 72 hours at 26°C, followed by an additional similar transfer and 48-hours incubation at 22°C. Subsequently, 5mL of cultures were transferred to 20 mL sterile seawater (SW) and incubated at 22°C for 24 hours in sterile glass tubes. SAR11 HTCC1062 was cultured individually in low-nutrient media supplemented with 80 µM pyruvate, 40 µM oxaloacetate, 40 µM taurine, or 50 µM glycine as the sole carbon source. Cultures were grown for 10 days until reaching the late exponential phase, indicating carbon source depletion. SAR11 cells were counted using Guava EasyCyte Plus flow cytometer, and all other cultures were counted using Cytoflex flow cytometer. Bacteria (final concentration of $10^5$, similar to seawater cell counts) were added to sterile SW (final volume of 80mL in 300mL bottles) supplemented with $NH_4$ and $PO_4$ and either nSM or vSM at microcosm-equivalent concentrations. Samples were incubated in darkness at 22°C for ~20 hr, after which samples for FCM analysis were collected, and radioactive incubations (4 hr) for BP and DCF measurements were initiated.

**Supplementary text 5 - Plaque assay for quantifying viable phages**

The plaque assay of phage P-HM2 was used to quantify viable phages for infecting *Prochlorococcus* MED4. *Alteromonas* DE served as a helper bacterium [18] for plating *Prochlorococcus* on low melting point (LMP) agarose (Invitrogen, UltraPure LMP agarose, cat no. 16520-100) plates. *Alteromonas* DE was prepared by centrifuging 1ml of overnight culture growing in ProMM [18] at 8000 RPM for 2 minutes, resuspending in 1ml of Pro99, and repeating the washing step twice before final resuspension in 1ml Pro99. All solutions were prepared using Milli-Q water (Millipore system). $Na_2SO_3$ stocks (1M concentration) was freshly prepared on the day of plating and sterilized using a 0.2µm syringe filter. A 0.35% LMP agarose solution was prepared by adding 0.35g of LMP agarose per 100ml of autoclaved MSW and microwaving to dissolve. After cooling to ~33ºC, nutrients were added to Pro99 medium (800µM $NH_4$, 50µM $PO_4$, 10,000X trace metals) supplemented with 1M $Na_2SO_3$. The agar solution was allowed to cool to ~30ºC before use. Phage lysate was serially diluted ($10^{-1}$ to $10^{-6}$) in Pro99 before plating. For plating, 2.7 mL of *Prochlorococcus* (~$10^8$ cells), 100µl of *Alteromonas* (~$10^6$ resuspension in Pro99) and 200µl of diluted lysate were added and mixed by pipetting. Then, 12ml of 0.35% LMP agarose was added to the opposite side of the plate, and the plates were mixed gently before agar was solidified. Plates were allowed to dry for 1 hour, sealed with parafilm, and incubated at ~22ºC on a net to prevent direct contact with the glass tray, reducing heating and steam accumulation. Plaques were typically observed within a week.

**Supplementary text 6 – Samples collection for the laboratory experiment with *Prochlorococcus***

Samples for protein, RNA and DNA quantification were collected at each timepoint in triplicate or duplicate (for proteins) by filtering ~50 mL on 0.7 µm GF/F, preserved in lysis buffer (40 mM EDTA, 50 mM Tris, pH 8.3, 0.75 M sucrose), and stored at -80˚C until further analysis. Flow cytometry samples were fixed with a final concentration of 0.125% glutaraldehyde for 10 minutes, frozen, and stored at -80˚C until counted using BD Canto II flow-cytometer. Additionally, samples for measuring $NH_4$, $PO_4$, DOC and dissolved protein, RNA and DNA (as described below) were taken from the media after filtration (0.2 μm polycarbonate filters) and stored at -20˚C. All remaining lysate from $T_2$ (infected) and $T_3$ (uninfected) was retained at -20˚C for the microcosm experiment.

**Supplementary text 7 – Measuring DOC and TON in spent-media**

DOC samples were collected into cleaned, pre-combusted, acid washed (10% HCL) 40ml vials, added with 20µL of 36% HCL to remove dissolved inorganic C before preservation at 4ºC under dark. DOC was measured a few days later via InnovOx Analyzer (Serial Number 1051, Version 03.10.017). TN samples were collected into cleaned 50ml falcon tubes and preserved at -20ºC until measured photometrically using a Foss Analytical FIAstar 5000 115 Analyzer with DDW detector and a common standard for TN (AN 5202D, DIN EN ISO 117 13395, ISO 11905). Dissolved organic nitrogen (DON) concentrations were calculated by subtracting ammonium concentrations from the total nitrogen values.

**Supplementary text 8 - 18S rRNA sequence analysis**

Sequences were processed using the DADA2 pipeline, following the same approach as for 16S. Forward and backward reads were truncated after quality inspections to 150 and 150 bases. After sequences merging, a consensus length of only between 125 and 175 bases was accepted. Finally, amplicon sequence variants (ASVs) that have less than 100 in total (all samples) were removed, remaining with a total of 368 sequences. Taxonomic classification was carried out in MEGAN6 (v6.24) using BLASTn results, excluding uncultured or environmental sequences. Assignments were made using the LCA algorithm with the MeganMapDB. Parameters included a Top Percent range of 3–10% and a Minimum Score threshold of 75–100. All bacteria, mitochondria, chloroplast, and ASVs without any taxonomic affiliation were discarded from downstream analyses. Full ASVs and taxonomy list of manually curated 18S rRNA can be found in Supplementary file 3.

**Supplementary text 9 – MAGs reconstruction**

For reconstructing metagenome assembled genomes (MAGs), assembled contigs for each biological condition were separately binned using MetaBat2 [19]. A quality assessment i.e. degree of genome completeness and contamination of the genome bins or MAGs was performed using CheckM [20]. Finally, we performed taxonomic classification of the high-quality MAGs (<10% contamination) by comparing with publicly available bacterial and archaeal genomes in the Genome Taxonomy Database (GTDB) using the tool GTDB-tk [21]. Most of the MAGs had >85% completion (70 MAGs), but few (13 MAGs) had 70-85% completion in an attempt to include more taxa identified in the 16S sequencing. The completeness of metabolic pathways within their genomes was performed via KEGG decoder script [22], in which ‘anaplerotic genes’ includes MD, PEPC, PEPCK and PC. Full list of software and their version is available on Table S2. For detailed MAGs list and assigned KEGGs see Supplementary file 9.

**Phylogenomic tree.** A phylogenomic tree was constructed using Anvi'o (v8) [23] based on eight universally conserved ribosomal protein genes (*Ribosomal_L1, L2, L3, L4, L5, L6, S7,* and *S8*). For each genome, the best-matching amino acid sequence of each target gene was identified and extracted. The resulting protein sequences were aligned, concatenated, and used to infer a maximum likelihood phylogeny with FastTree2 under default parameters. The final tree was visualized using Anvi'o's interactive interface. See Table S1 for details on the bacterial genomes.

### References


1. Reich T, Ben-Ezra T, Belkin N *et al.* A year in the life of the Eastern Mediterranean: Monthly dynamics of phytoplankton and bacterioplankton in an ultra-oligotrophic sea. *Deep Sea Res Part I Oceanogr Res Pap* 2022;**182**:103720.

2. Reich T, Belkin N, Sisma-Ventura G *et al.* Significant dark inorganic carbon fixation in the euphotic zone of an oligotrophic sea. *Limnol Oceanogr* 2024;**69**:1129–42.

3. Braun A, Spona-Friedl M, Avramov M *et al.* Reviews and syntheses: Heterotrophic fixation of inorganic carbon – significant but invisible flux in environmental carbon cycling. *Biogeosciences* 2021;**18**:3689–700.

4. Signori CN, Valentin JL, Pollery RCG *et al.* Temporal Variability of Dark Carbon Fixation and Bacterial Production and Their Relation with Environmental Factors in a Tropical Estuarine System. *Estuaries and Coasts* 2018;**41**:1089–101.

5. Givati S, Forchielli E, Aharonovich D *et al.* Diversity in the utilization of different molecular classes of dissolved organic matter by heterotrophic marine bacteria. Biddle JF (ed.). *Appl Environ Microbiol* 2024;**90**:e00256-24.

6. Haider MN, Iqbal MM, Nishimura M *et al.* Bacterial response to glucose addition: growth and community structure in seawater microcosms from North Pacific Ocean. *Sci Rep* 2023;**13**:341.

7. Eilers H, Pernthaler J, Amann R. Succession of Pelagic Marine Bacteria during Enrichment: a Close Look at Cultivation-Induced Shifts. *Appl Environ Microbiol* 2000;**66**:4634–40.

8. Baltar F, Lundin D, Palovaara J *et al.* Prokaryotic Responses to Ammonium and Organic Carbon Reveal Alternative CO2 Fixation Pathways and Importance of Alkaline Phosphatase in the Mesopelagic North Atlantic. *Front Microbiol* 2016;**7**:1–19.

9. Eigemann F, Rahav E, Grossart H *et al.* Phytoplankton exudates provide full nutrition to a subset of accompanying heterotrophic bacteria via carbon, nitrogen and phosphorus allocation. *Environ Microbiol* 2022;**24**:2467–83.

10. Pinhassi J, Sala MM, Havskum H *et al.* Changes in Bacterioplankton Composition under Different Phytoplankton Regimens. *Appl Environ Microbiol* 2004;**70**:6753–66.

11. Riemann L, Steward GF, Azam F. Dynamics of Bacterial Community Composition and Activity during a Mesocosm Diatom Bloom. *Appl Environ Microbiol* 2000;**66**:2282–2282.

12. Eigemann F, Rahav E, Grossart HP *et al.* Phytoplankton Producer Species and Transformation of Released Compounds over Time Define Bacterial Communities following Phytoplankton Dissolved Organic Matter Pulses. *Appl Environ Microbiol* 2023;**89**, DOI: 10.1128/aem.00539-23.

13. Xiao X, Guo W, Li X *et al.* Viral Lysis Alters the Optical Properties and Biological Availability of Dissolved Organic Matter Derived from Prochlorococcus Picocyanobacteria. Johnson KN (ed.). *Appl Environ Microbiol* 2021;**87**:1–19.

14. Henshaw RJ, Moon J, Stehnach MR *et al.* Early viral infection of cyanobacteria drives bacterial chemotaxis in the oceans. *bioRxiv* 2023:2023.10.24.563588.

15. Teeling H, Fuchs BM, Becher D *et al.* Substrate-Controlled Succession of Marine Bacterioplankton Populations Induced by a Phytoplankton Bloom. *Science (80- )* 2012;**336**:608–11.

16. Buchan A, LeCleir GR, Gulvik CA *et al.* Master recyclers: features and functions of bacteria associated with phytoplankton blooms. *Nat Rev Microbiol* 2014;**12**:686–98.

17. Ahn S, Jung J, Jang I-A *et al.* Role of Glyoxylate Shunt in Oxidative Stress Response. *J Biol Chem* 2016;**291**:11928–38.

18. Morris JJ, Kirkegaard R, Szul MJ *et al.* Facilitation of robust growth of Prochlorococcus colonies and dilute liquid cultures by "helper" heterotrophic bacteria. *Appl Environ Microbiol* 2008;**74**:4530–4.

19. Kang DD, Li F, Kirton E *et al.* MetaBAT 2: an adaptive binning algorithm for robust and efficient genome reconstruction from metagenome assemblies. *PeerJ* 2019;**7**:e7359.

20. Parks DH, Imelfort M, Skennerton CT *et al.* CheckM: assessing the quality of microbial genomes recovered from isolates, single cells, and metagenomes. *Genome Res* 2015;**25**:1043–55.

21. Chaumeil P-A, Mussig AJ, Hugenholtz P *et al.* GTDB-Tk: a toolkit to classify genomes with the Genome Taxonomy Database. Hancock J (ed.). *Bioinformatics* 2020;**36**:1925–7.

22. Graham ED, Heidelberg JF, Tully BJ. Potential for primary productivity in a globally-distributed bacterial phototroph. *ISME J* 2018;**12**:1861–6.

23. Eren AM, Esen ÖC, Quince C *et al.* Anvi'o: an advanced analysis and visualization platform for 'omics data. *PeerJ* 2015;**3**:e1319.

### Supplementary Tables

**Table S1. List of strains used in the cultures experiments (Figure 5).**

| Strain | Genus | NCBI taxonID | Length (Mbp) | GC % | Completeness % |
|---|---|---|---|---|---|
| **DE** | *Alteromonas mediterranea DE* | 1774373 | 4.5 | 44.9% | 99.8% |
| **TAC** | *Pseudoalteromonas haloplanktis* | 326442 | 3.9 | 40.1% | 100.0% |
| **MMB-1** | *Marinomonas mediterranea* | 717774 | 4.7 | 44.1% | 99.9% |
| **KMM** | *Formosa agariphila* | 1347342 | 4.2 | 33.3% | 99.0% |
| **HTCC1062** | *Candidatus Pelagibacter ubique* | 335992 | 1.3 | 29.5% | 99.9% |
| **HOT5_F3** | *Rhodobacter marinovum* | 1291154 | | | |

**Table S2. List of software versions used for metagenomic analysis.**

| Program | Version | Relevant Links |
|---|---|---|
| **FastQC** | 0.11.9 | https://www.bioinformatics.babraham.ac.uk/projects/fastqc/ |
| **MultiQC** | 1.11 | https://multiqc.info/ |
| **bbduk** | 38.86 | https://jgi.doe.gov/data-and-tools/software-tools/bbtools/bb-tools-user-guide/ |
| **megahit** | 1.2.9 | https://github.com/voutcn/megahit#megahit |
| **bit** | 1.8.53 | https://github.com/AstrobioMike/bioinf_tools#bioinformatics-tools-bit |
| **bowtie2** | 2.3.5.1 | https://github.com/BenLangmead/bowtie2#overview |
| **samtools** | 1.9 | https://github.com/samtools/samtools#samtools |
| **prodigal** | 2.6.3 | https://github.com/hyattpd/Prodigal#prodigal |
| **KOFamScan** | 1.3.0 | https://github.com/takaram/kofam_scan#kofamscan |
| **CAT** | 5.2.2 | https://github.com/dutilh/CAT#cat-and-bat |
| **Metabat2** | 2.15 | https://bitbucket.org/berkeleylab/metabat/src/master/ |
| **checkm** | 1.1.3 | https://github.com/Ecogenomics/CheckM |
| **gtdbtk** | 1.5.0 | https://github.com/Ecogenomics/GTDBTk |
| **KEGGDecoder** | 1.2.2 | https://github.com/bjtully/BioData/tree/master/KEGGDecoder#kegg-decoder |
| **HUMAnN3** | 3.6 | https://huttenhower.sph.harvard.edu/humann3/ |
| **MetaPhlAn3** | 4.0.1 | https://github.com/biobakery/MetaPhlAn/tree/3.0 |

### Supplementary Figures

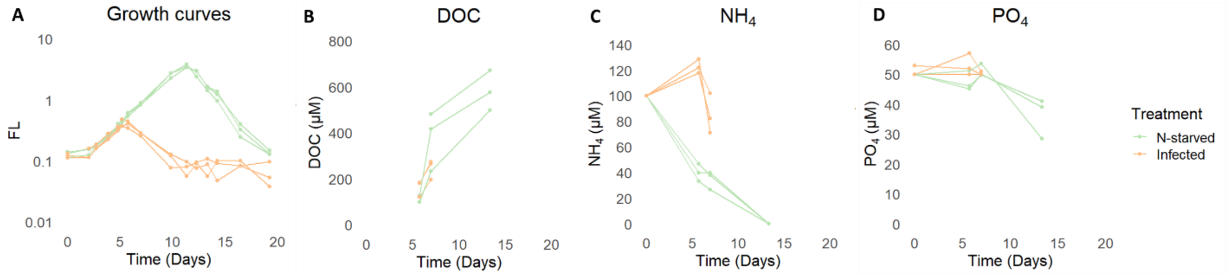


**Figure S1**. **Changes in nutrients and dissolved organic carbon (DOC) concentrations along *Prochlorococcus* growth curve**.

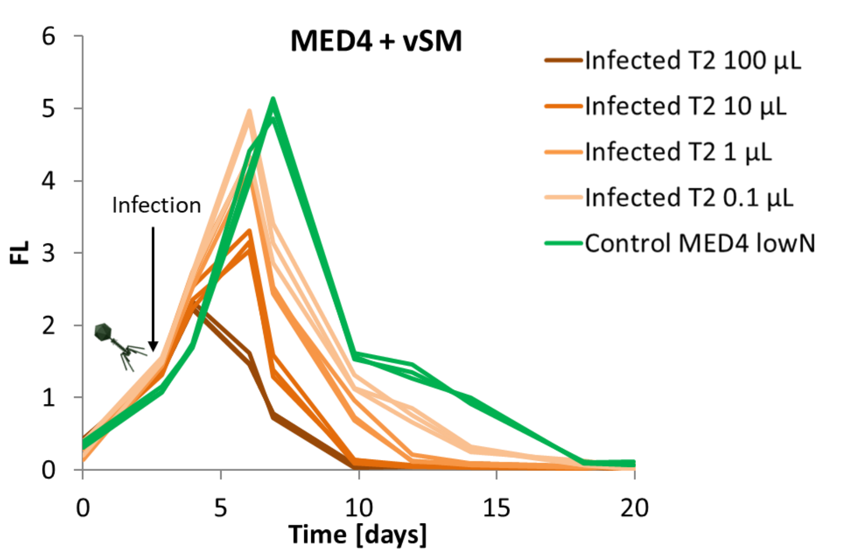


**Figure S2**. ***Prochlorococcus* MED4 infection following addition of experimental lysate**. The highest volume added (100 µL) caused the most pronounced lysis, with all lysate treatments showing significant differences compared to the control (MED4 in lowN media). The infection occurred on day 3.

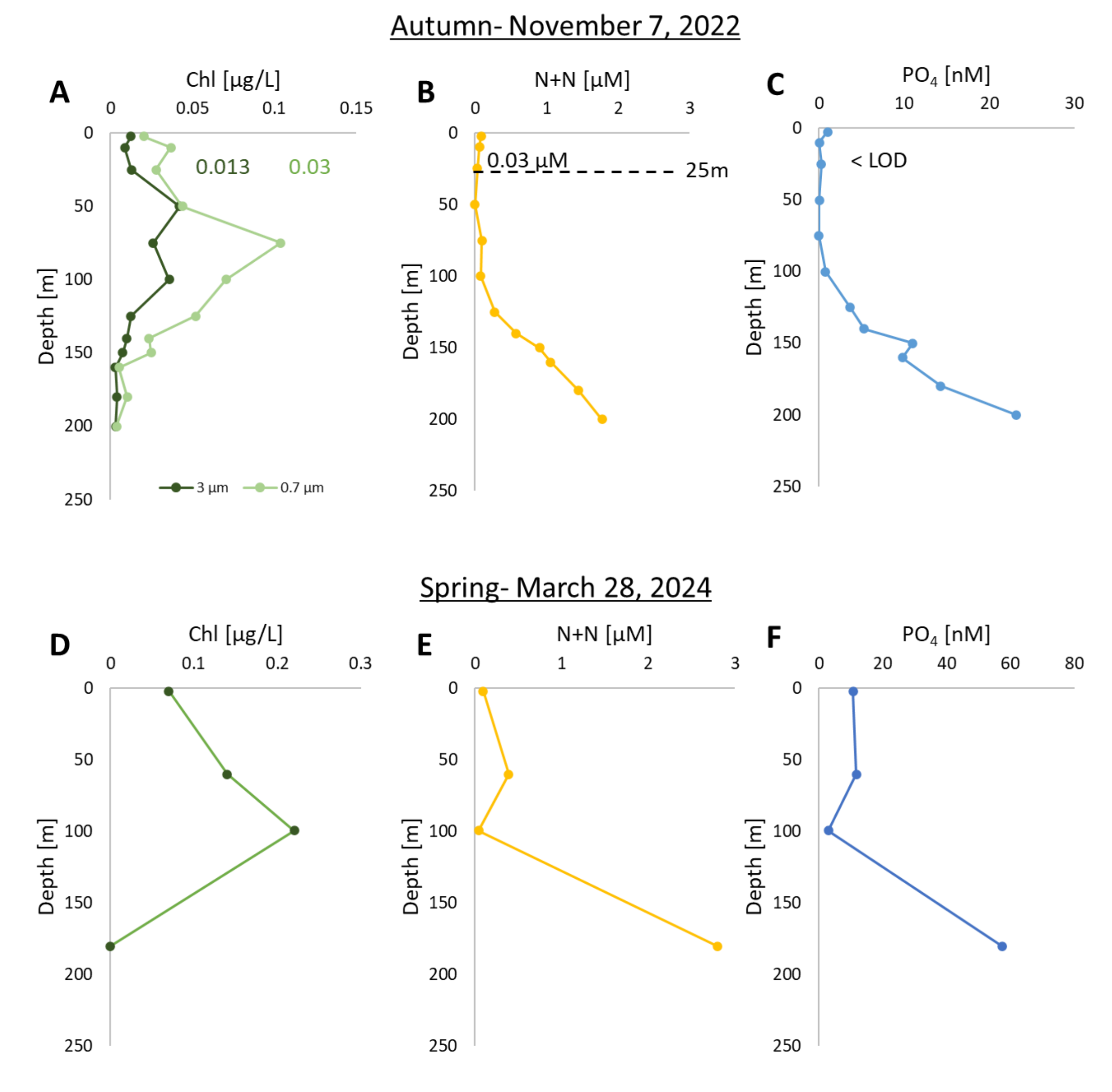


**Figure S3**. **Water profiles from Nov 7, 2022**. (A) Chlorophyll-*a* concentration (µg/L) measured using chlorophyll extraction for large phytoplankton (>3µm) and small cyanobacteria (0.7-3µm). (B) Concentration of dissolved inorganic nitrogen ($NO_2$+$NO_3$) (µM) (C) Concentrations of dissolved inorganic $PO_4$ (nM) were below limit of detection (LOD = 6nM) in the upper water column. Measurements were taken by Alon Alon Blachinsky from the Morris Kahn Research Station, Charney School of Marine Sciences. (D) Chlorophyll-*a* concentration (µg/L) measured using chlorophyll extraction for total phytoplankton community. (E) Concentration of $NO_2$+$NO_3$ (µM). (F) Concentration of dissolved inorganic $PO_4$ (nM). Measurements were taken by the IOLR during a transect cruise (H51). LOD = limit of detection. Concentration of nutrients were 0.03 µM for total N ($NO_2$ and $NO_3$) and below limit of detection for phosphate (Figure S3B, C). Depth of the deep chlorophyll maximum (DCM) was ~75m and was mostly comprised of small phytoplankton fraction, as suggested by the fractionated chlorophyll extraction (Figure S3A). In the spring experiment, concentration of chlorophyll (approximately 0.1 µg/L) and nutrients (approximately 10 nM for $PO_4$ and ~0.25 µM for $NO_2$ and $NO_3$) were higher than November (Figure S3D-F), consistent with expected seasonal patterns.

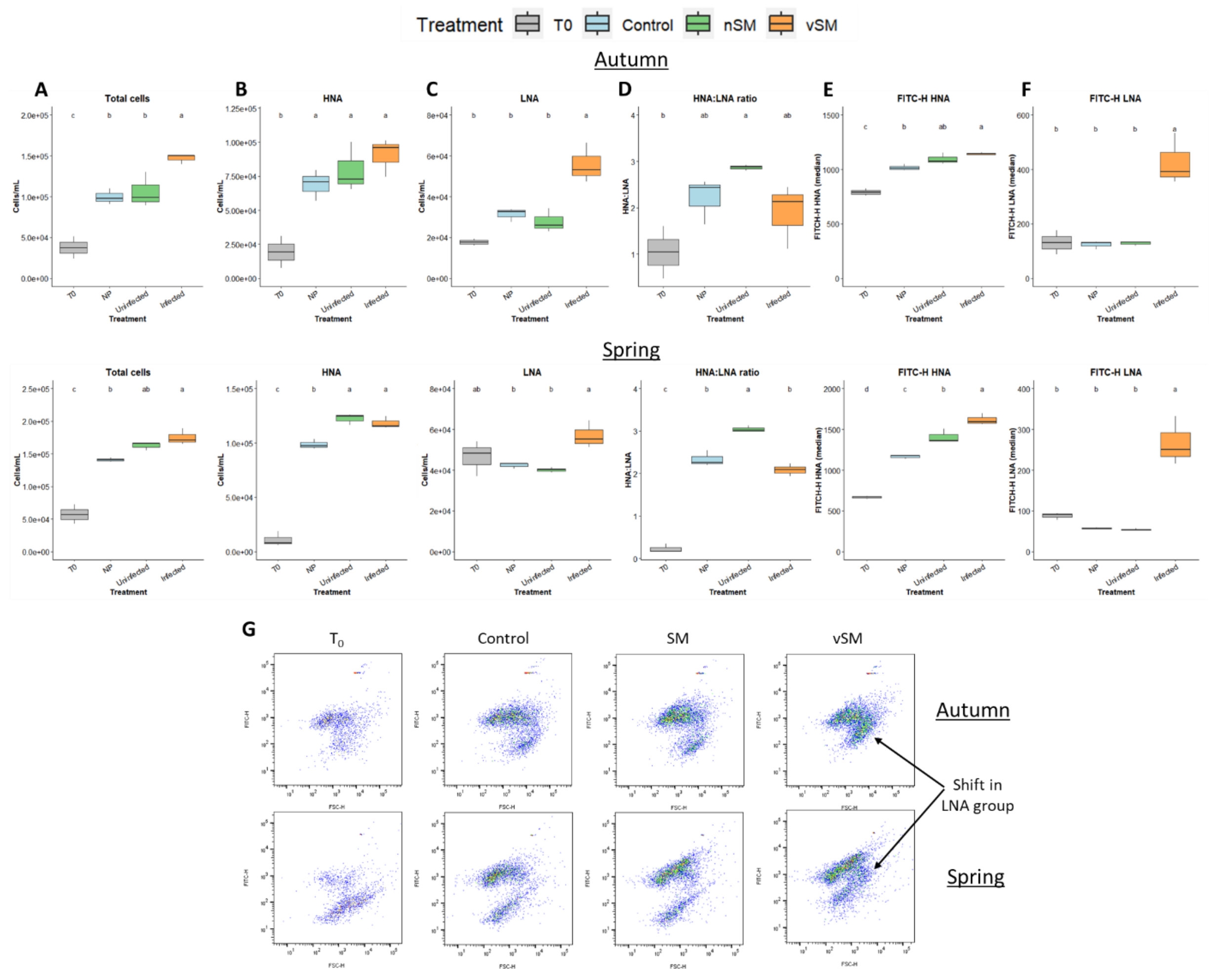


**Figure S4**. **Spent media from N-starved and infected *Prochlorococcus* differently affect microbial counts by flow cytometry.** (A) Total cells count. (B) High nucleic acids population (HNA) (cells/mL). (C) Low nucleic acids population (LNA) (cells/mL). (D) The ratio between HNA:LNA, potential indicator for higher activity. (E) Median fluorescein isothiocyanate (FITC) for the HNA. (F) Median fluorescein isothiocyanate (FITC) for the LNA. (G) Scattergram showing the shift in LNA acid with the addition of vSM. The different letters above the box plots indicate statistically significant differences among the treatments (one-way ANOVA and post-hoc Tukey test, $p < 0.05$).

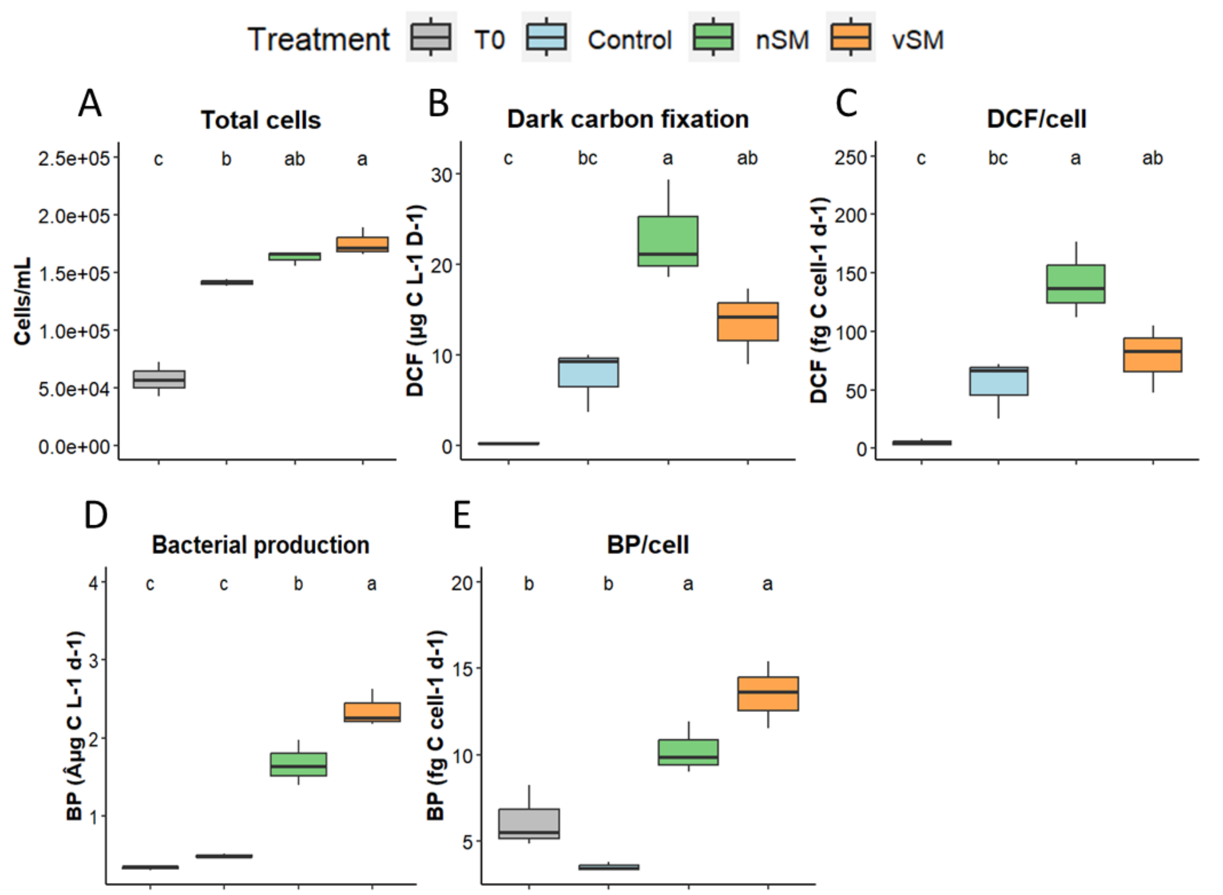


**Figure S5**. **Spent media from N-starved and infected *Prochlorococcus* differently affect natural microbial community activity - spring.** (A) Total cells count. (B) Dark inorganic carbon fixation (DCF). (C) Per-cell DCF. (D) Bacterial productivity (BP). (E) Per-cell BP. Box-Whisker plots show the interquartile range (25th–75th percentile) of the data set. The horizontal line within the box represents the median value (N=3). The different letters above the box plot indicate statistically significant differences among the treatments (one-way ANOVA and post-hoc Tukey test, $p < 0.05$).

229

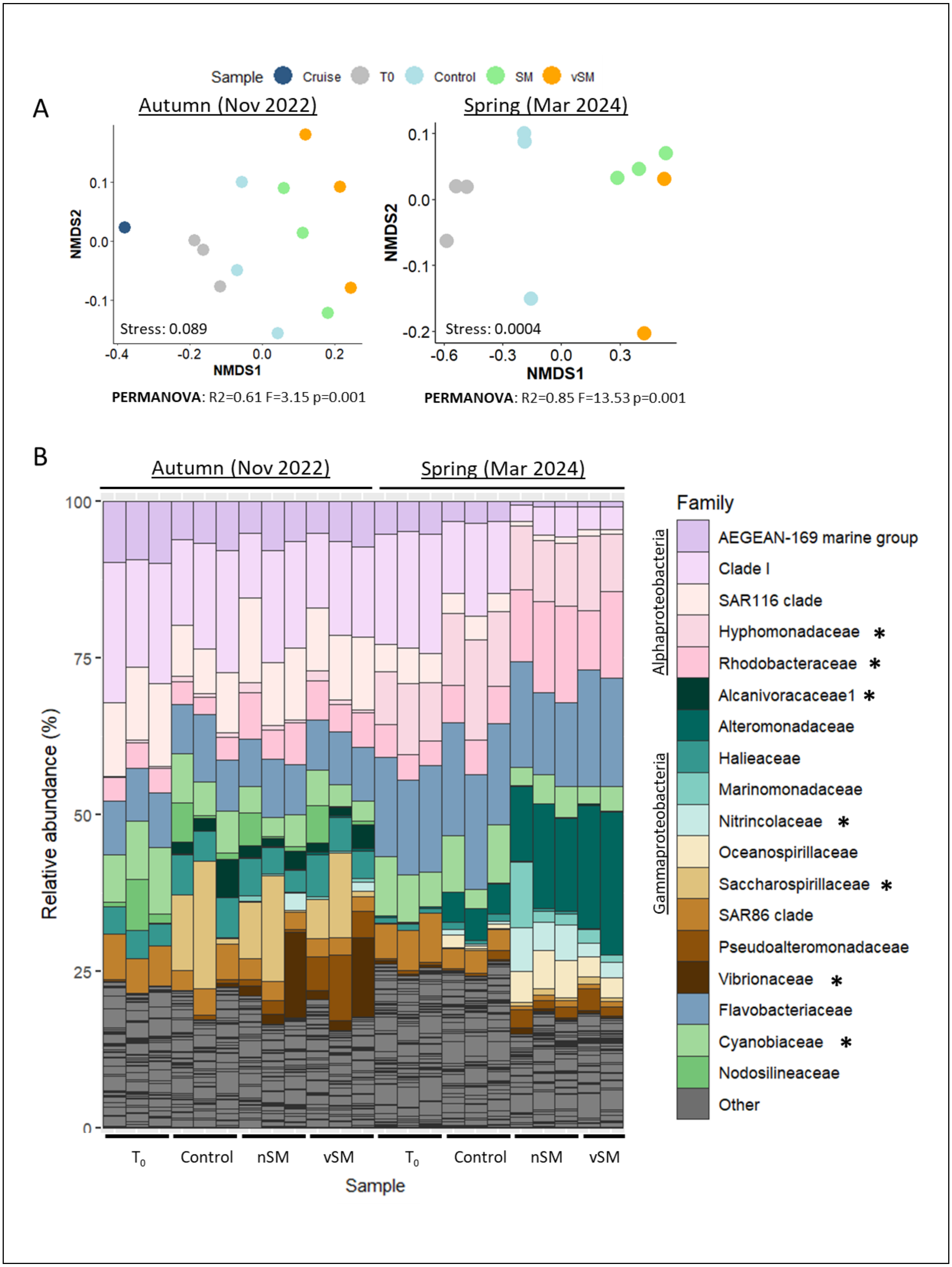

Sample
Cruise
T0
Control
SM
vSM
A
Autumn (Nov 2022)
Spring (Mar 2024)
NMDS2
NMDS1
Stress: 0.089
Stress: 0.0004
PERMANOVA: R2=0.61 F=3.15 p=0.001
PERMANOVA: R2=0.85 F=13.53 p=0.001
B
Autumn (Nov 2022)
Spring (Mar 2024)
Relative abundance (%)
Family
Alphaproteobacteria
Gammaproteobacteria
AEGEAN-169 marine group
Clade I
SAR116 clade
Hyphomonadaceae *
Rhodobacteraceae *
Alcanivoracaceae1 *
Alteromonadaceae
Halieaceae
Marinomonadaceae
Nitrincolaceae *
Oceanospirillaceae
Saccharospirillaceae *
SAR86 clade
Pseudoalteromonadaceae
Vibrionaceae *
Flavobacteriaceae
Cyanobiaceae *
Nodosilineaceae
Other
T0
Control
nSM
vSM
Sample

**Figure S6. Spent media from N-starved and infected *Prochlorococcus* effect on natural microbial community structure.** (A) NMDS plots based on Bray-Curtis dissimilarity for 16S amplicon sequencing in the Autum and Spring. Spring NMDS – 3 dimensions. Results of the PERMANOVA test are shown under both plots. (B) 16S rRNA stacked bar charts showing relative proportions (percentages of 16S rRNA sequences) of microbial community at the family level. Other taxa=families not greater than 5% in any sample. Families for which MAGs are available are marked with an asterisk.

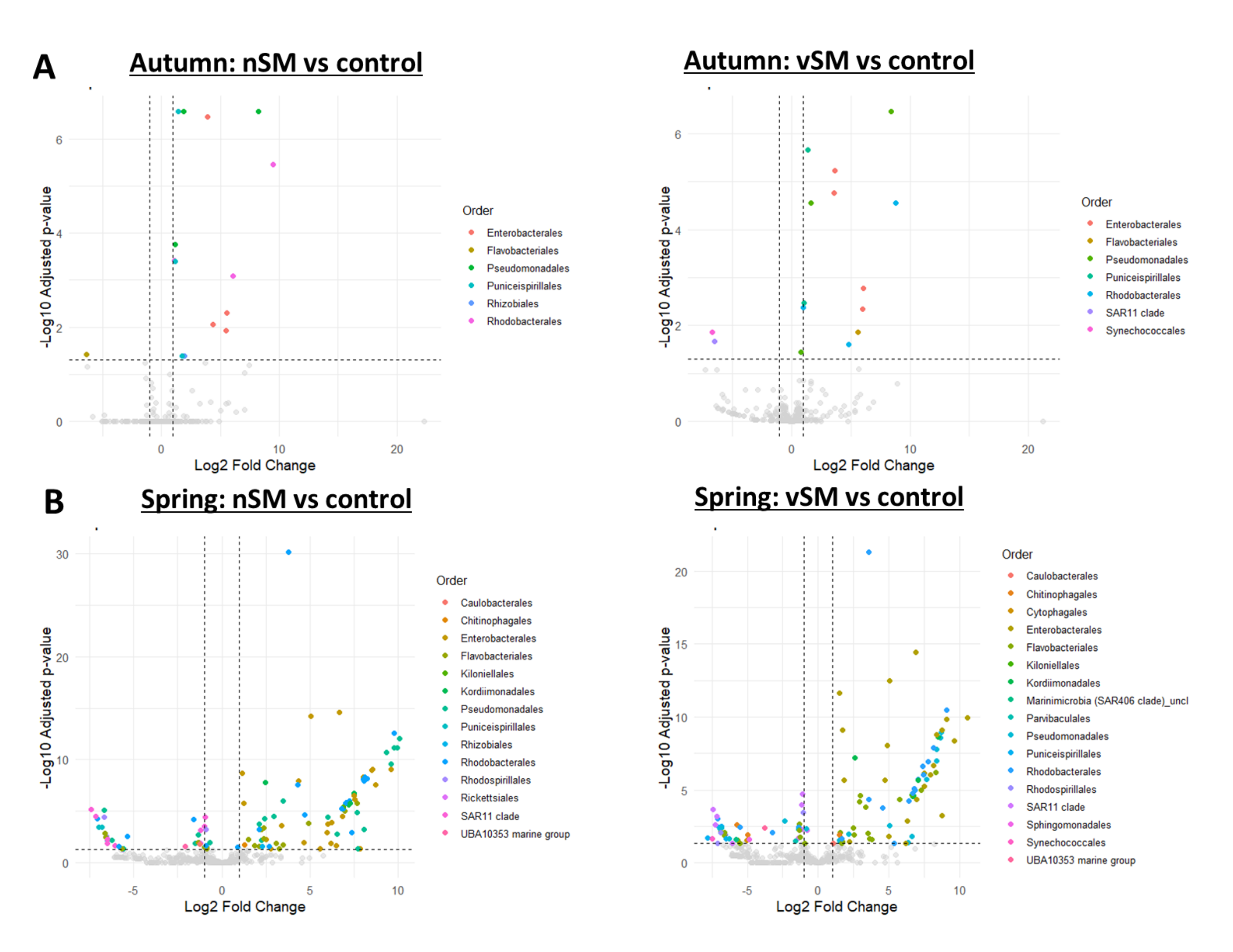


**Figure S7. DESeq analysis of 16S rRNA amplicon sequencing data at the order level from the autumn (A) and spring (B) experiments**. Both experiments and treatments show a significant increase in specific bacterial orders, including Enterobacterales (e.g. *Vibrionaceae, Pseudoalteromonadaceae, Alteromonadaceae*), Rhodobacterales (*Rhodobacteraceae*) and Pseudomonadales (e.g. *Marinomonadaceae, Nitrincolaceae*).

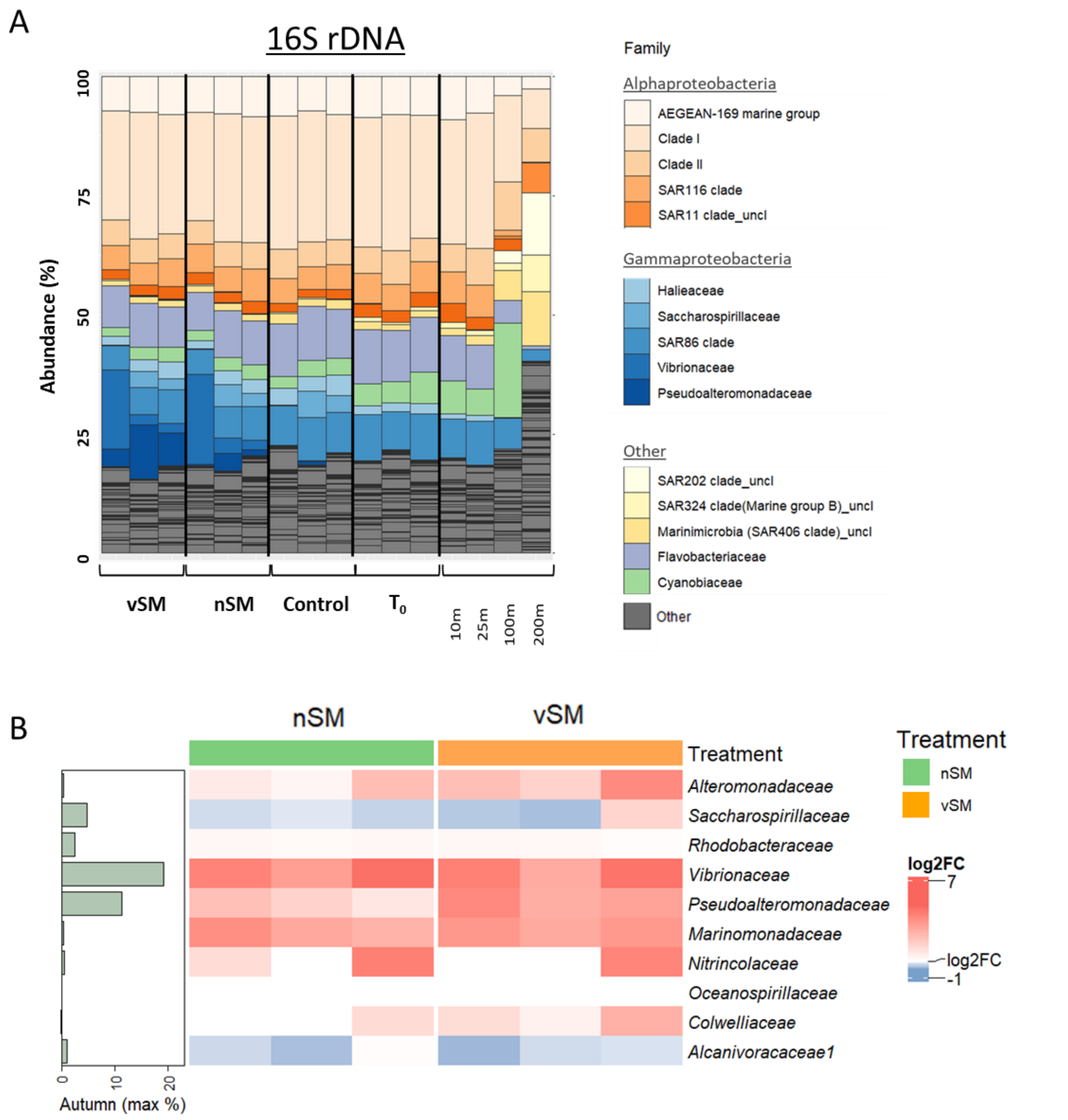


**Figure S8**. **16S rDNA amplicon sequencings (autumn, 2022)**. (A) 16S rDNA stacked bar charts showing relative proportions (percentages of 16S rDNA sequences) of microbial community at the family level. Other taxa=families not greater than 5% in any sample. (B) Heatmap showing top families corresponding the Figure 2D. Left side bar-plots showing maximum relative abundance in nSM or vSM in the autumn, based on (A).

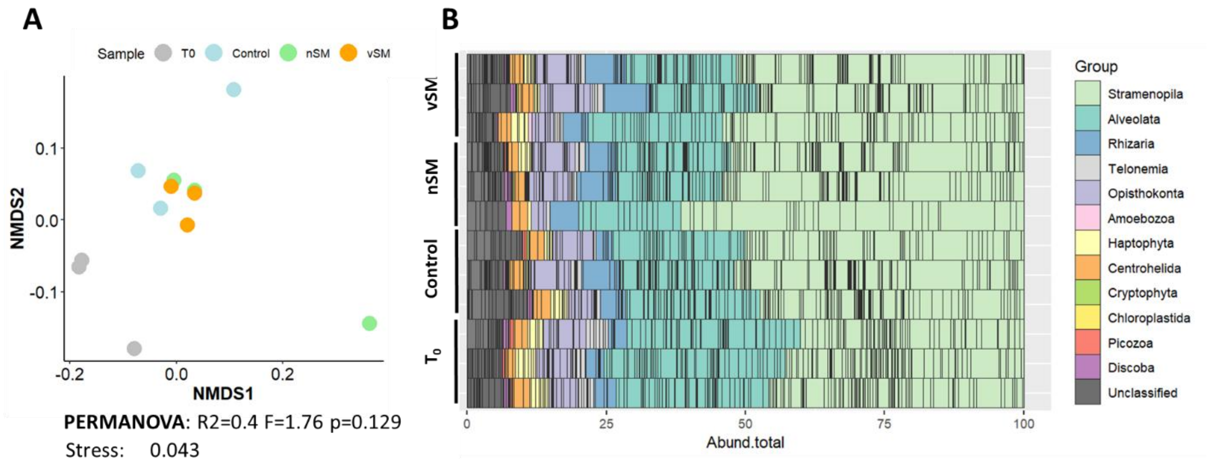


**Figure S9**. **Spent media from N-starved and infected *Prochlorococcus* effect on natural microbial community structure in the autumn experiment (18S rRNA).** (A) NMDS plots based on Bray-Curtis dissimilarity for 18S amplicon sequencing. Results of the PERMANOVA test are shown under the plot. (B) 168 rRNA stacked bar charts showing relative proportions (percentages of 16S rRNA sequences) of microbial community at the family level. Other groups not greater than 1% in any sample. No differences in community structure were observed between treatments and control, which slightly differ from $T_0$. Abbreviation: nSM- spent media following nitrogen starvation; vSM - viral spent media.

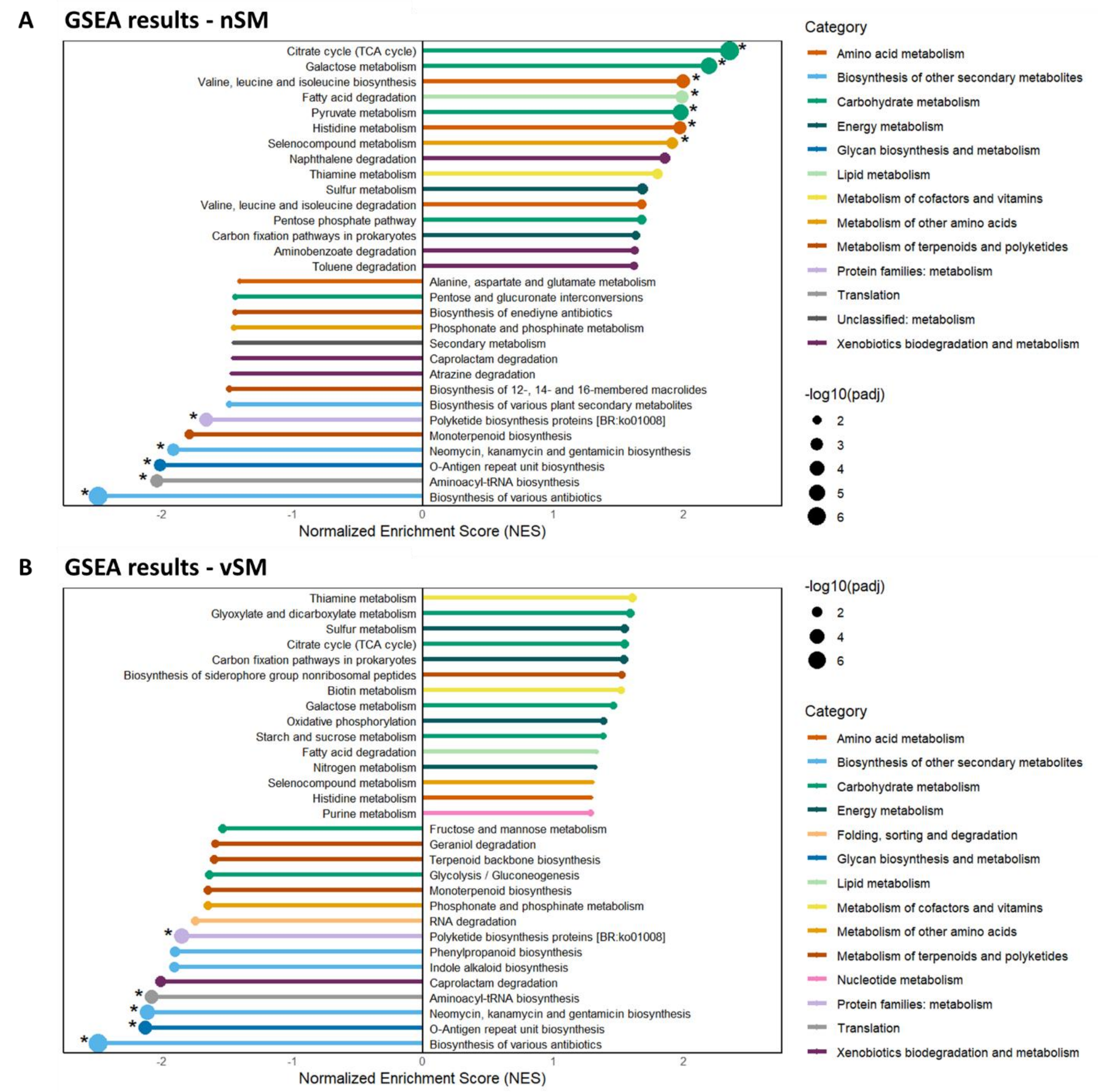


**Figure S10**. **GSEA results - top enriched pathways in nSM (up) and vSM (bottom)**. Pathways are colored by their KEGG categories. The size of the bars represents the normalized enrichment score (NES). The size of the points represents the adjusted *p*-value. Asterisk denotes statistically significant pathways (adjusted *p*-value < 0.05). *P*-value for individual KOs abundance was estimated using permutation test with 100000 permutations.

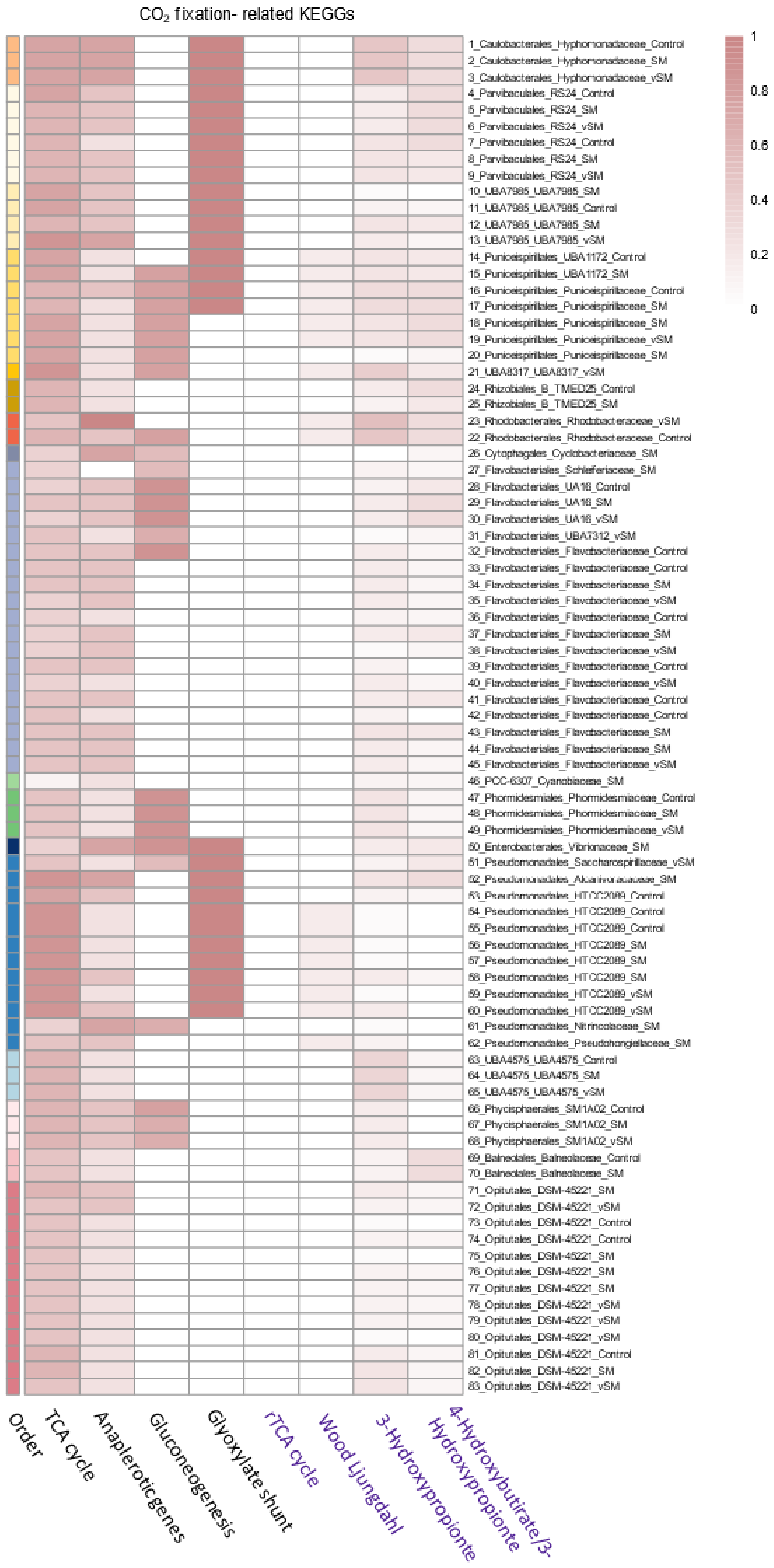
CO2 fixation- related KEGGs
1_Caulobacterales_Hyphomonadaceae_Control
2_Caulobacterales_Hyphomonadaceae_SM
3_Caulobacterales_Hyphomonadaceae_vSM
4_Parvibaculales_RS24_Control
5_Parvibaculales_RS24_SM
6_Parvibaculales_RS24_vSM
7_Parvibaculales_RS24_Control
8_Parvibaculales_RS24_SM
9_Parvibaculales_RS24_vSM
10_UBA7985_UBA7985_SM
11_UBA7985_UBA7985_Control
12_UBA7985_UBA7985_SM
13_UBA7985_UBA7985_vSM
14_Puniceispirillales_UBA1172_Control
15_Puniceispirillales_UBA1172_SM
16_Puniceispirillales_Puniceispirillaceae_Control
17_Puniceispirillales_Puniceispirillaceae_SM
18_Puniceispirillales_Puniceispirillaceae_SM
19_Puniceispirillales_Puniceispirillaceae_vSM
20_Puniceispirillales_Puniceispirillaceae_SM
21_UBA8317_UBA8317_vSM
24_Rhizobiales_B_TMED25_Control
25_Rhizobiales_B_TMED25_SM
23_Rhodobacterales_Rhodobacteraceae_vSM
22_Rhodobacterales_Rhodobacteraceae_Control
26_Cytophagales_Cyclobacteriaceae_SM
27_Flavobacteriales_Schleiferiaceae_SM
28_Flavobacteriales_UA16_Control
29_Flavobacteriales_UA16_SM
30_Flavobacteriales_UA16_vSM
31_Flavobacteriales_UBA7312_vSM
32_Flavobacteriales_Flavobacteriaceae_Control
33_Flavobacteriales_Flavobacteriaceae_Control
34_Flavobacteriales_Flavobacteriaceae_SM
35_Flavobacteriales_Flavobacteriaceae_vSM
36_Flavobacteriales_Flavobacteriaceae_Control
37_Flavobacteriales_Flavobacteriaceae_SM
38_Flavobacteriales_Flavobacteriaceae_vSM
39_Flavobacteriales_Flavobacteriaceae_Control
40_Flavobacteriales_Flavobacteriaceae_vSM
41_Flavobacteriales_Flavobacteriaceae_Control
42_Flavobacteriales_Flavobacteriaceae_Control
43_Flavobacteriales_Flavobacteriaceae_SM
44_Flavobacteriales_Flavobacteriaceae_SM
45_Flavobacteriales_Flavobacteriaceae_vSM
46_PCC-6307_Cyanobiaceae_SM
47_Phormidesmiales_Phormidesmiaceae_Control
48_Phormidesmiales_Phormidesmiaceae_SM
49_Phormidesmiales_Phormidesmiaceae_vSM
50_Enterobacterales_Vibrionaceae_SM
51_Pseudomonadales_Saccharospirillaceae_vSM
52_Pseudomonadales_Alcanivoracaceae_SM
53_Pseudomonadales_HTCC2089_Control
54_Pseudomonadales_HTCC2089_Control
55_Pseudomonadales_HTCC2089_Control
56_Pseudomonadales_HTCC2089_SM
57_Pseudomonadales_HTCC2089_SM
58_Pseudomonadales_HTCC2089_SM
59_Pseudomonadales_HTCC2089_vSM
60_Pseudomonadales_HTCC2089_vSM
61_Pseudomonadales_Nitrincolaceae_SM
62_Pseudomonadales_Pseudohongiellaceae_SM
63_UBA4575_UBA4575_Control
64_UBA4575_UBA4575_SM
65_UBA4575_UBA4575_vSM
66_Phycisphaerales_SM1A02_Control
67_Phycisphaerales_SM1A02_SM
68_Phycisphaerales_SM1A02_vSM
69_Balneolales_Balneolaceae_Control
70_Balneolales_Balneolaceae_SM
71_Opitutales_DSM-45221_SM
72_Opitutales_DSM-45221_vSM
73_Opitutales_DSM-45221_Control
74_Opitutales_DSM-45221_Control
75_Opitutales_DSM-45221_SM
76_Opitutales_DSM-45221_SM
77_Opitutales_DSM-45221_SM
78_Opitutales_DSM-45221_vSM
79_Opitutales_DSM-45221_vSM
80_Opitutales_DSM-45221_vSM
81_Opitutales_DSM-45221_Control
82_Opitutales_DSM-45221_SM
83_Opitutales_DSM-45221_vSM
1
0.8
0.6
0.4
0.2
0
Order
TCA cycle
Anapleroticgenes
Gluconeogenesis
Glyoxylate shunt
rTCA cycle
Wood Ljungdahl
3-Hydroxypropionte
4-Hydroxybutirate/3-Hydroxypropionte

**Figure S11**. **Heatmap showing distribution of pathways related to $CO_2$ fixation across the MAGs**. Purple columns names – autotorphic $CO_2$ fixation pathways. Black columns names – anaplerotic and related pathways. The analysis MAGs revealed that none harbored complete KEGG pathways for autotrophic CO2 fixation. KEGG annotations for the reductive tricarboxylic acid (rTCA) cycle were absent in all MAGs. While KEGG annotations for the Wood-Ljungdahl and 4-Hydroxybutyrate/3-Hydroxypropionate pathways were detected in several MAGs, their completeness remained low (<30%). The 3-Hydroxypropionate Bicycle exhibited the highest overall completeness, though most MAGs remained below 40%, with the highest completeness was observed in *Rhodobacteraceae* MED-G52 from the vSM samples (53%).

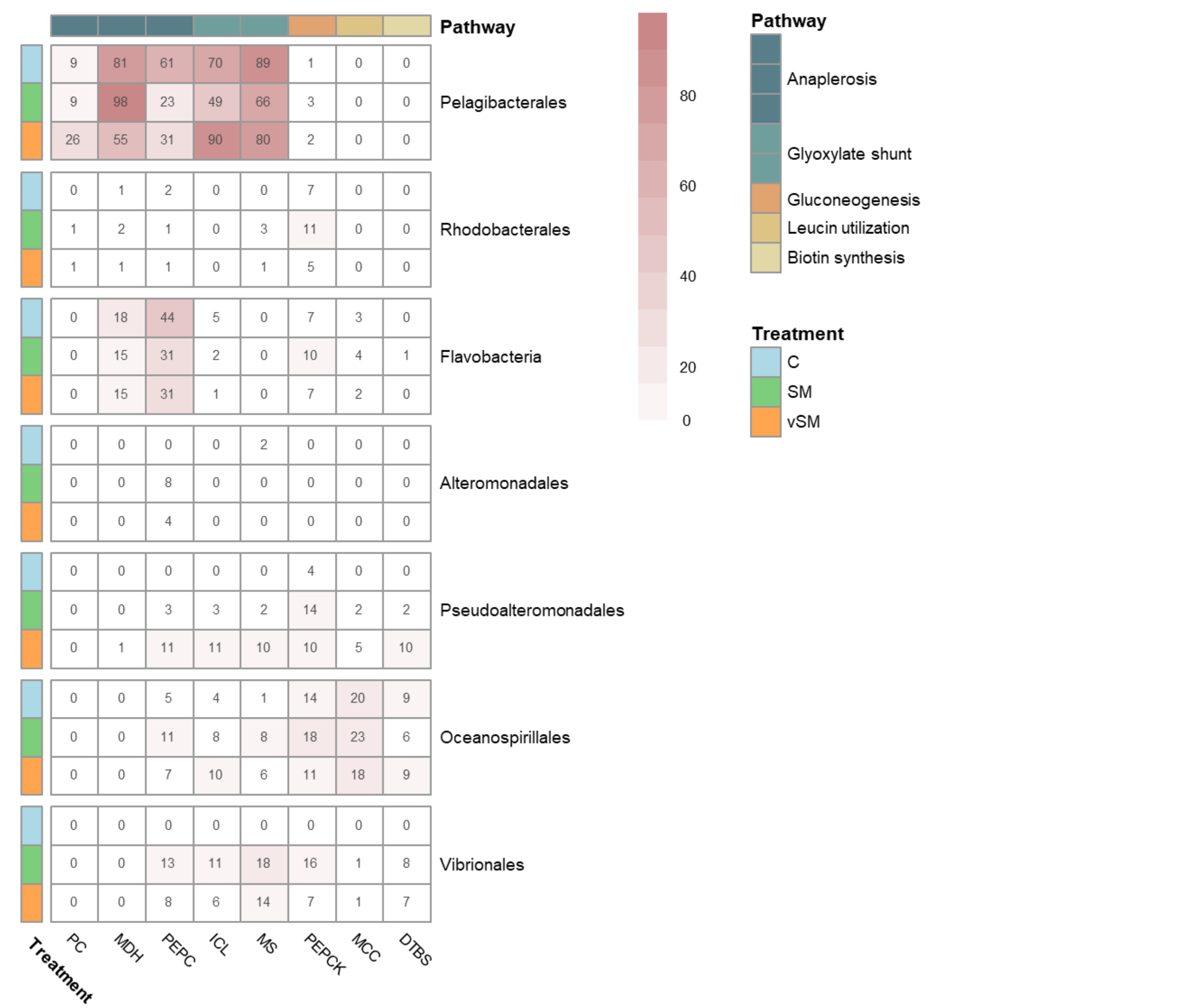


**Figure S12. Heatmap for average read coverage (CPM, N=3) for major heterotrophic taxa which exhibited the highest relative abundance of genes from Figure 4A.**

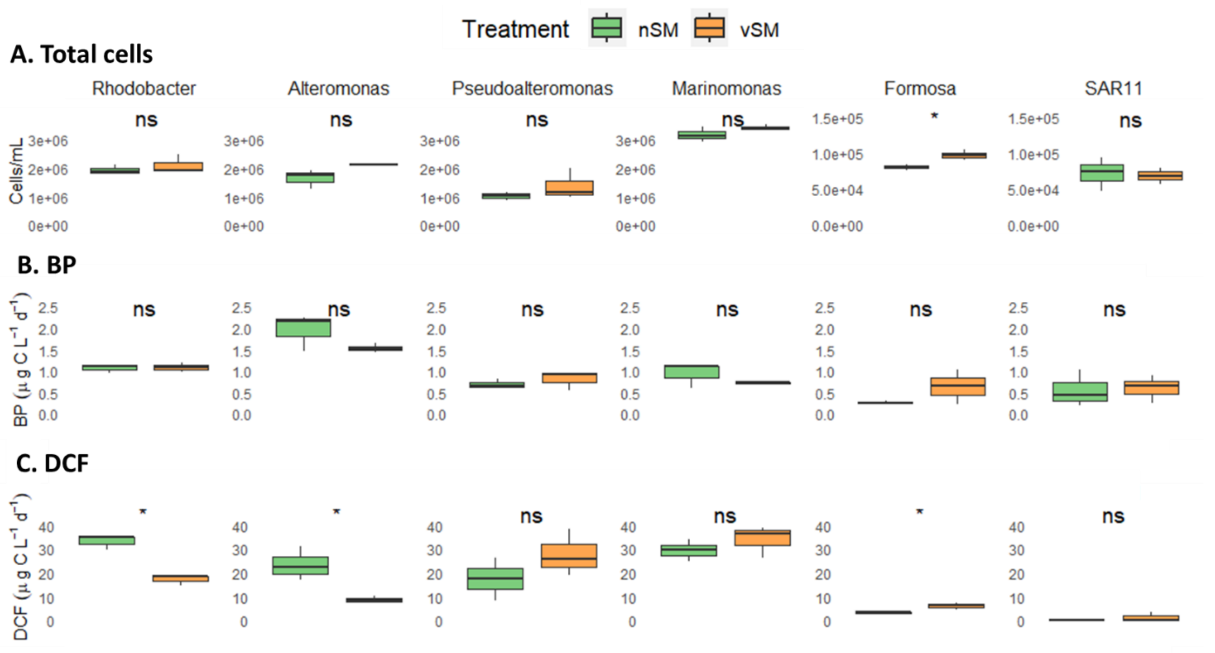


**Figure S13. The impact of spent media from N-starved (nSM) and infected (vSM) *Prochlorococcus* on BP and DCF rates in six C-starved laboratory cultures.** (A) Bacterial production (BP). (B) Dark inorganic carbon fixation (DCF). Box-Whisker plots show the interquartle range (25th–75th percentile) of the data set. The horizontal line within the box represents the median value (N=3). An asterisk denotes statistically significant results, determined by a paired t-test with Bonferroni correction. Note the different Y-axis between Formosa and SAR11 and the other strains in the total cells count, which may be due to technical issues during flow cytometry. *Rhodobacter – Roseobacteraceae Marinovum* HOT5_F3 (order of *Rhodobacterales*); *Alteromonas – Alteromonas mediterranea* DE; *Pseudoalteromonas – Pseudoalteromonas haloplanktis* TAC; *Marinomonas – Marinomonas mediterranea* MMB1; *Formosa – Formosa agariphila* KMM (Family of *Flavobacteriaceae*); SAR11 - Candidatus pelagibacteraceae St. HTCC1062 (Table S1).
